# Marfan Patient iPSC-Derived Endothelial Cells Carrying *FBN1* Variants Reveal Endothelial Dysfunction

**DOI:** 10.64898/2026.08.20.745919

**Authors:** Philipp C. Hauger, Gabija Danilinaite, Leonardo Spagnolello, Carsten Künne, Max C. Overboom, Jan Willem Buikema, Vivian de Waard, Peter L. Hordijk

## Abstract

Marfan syndrome (MFS) is an inherited connective tissue disorder caused by pathogenic variants in FBN1, encoding fibrillin-1, with life-threatening aortic complications arising in part from endothelial cell (EC) dysfunction. To study this in a human model, we generated hiPSC-derived ECs from three MFS patients (iMFS-ECs). We show that iMFS-ECs recapitulate known disease phenotypes, including impaired alignment in the direction of flow. Moreover, we found that iMFS-ECs do not recover from TNF-α-induced loss of barrier integrity, due to sustained EC contractility. iMFS-ECs exhibited TNF-α-induced ICAM1 upregulation and NF-κB activation comparable to healthy donor-derived hiPSC-ECs by bulk RNA-seq, while expression of genes linked to cytoskeletal arrangements, cell signaling and ECM remodeling were dysregulated. In conclusion, we show that hiPSC derived ECs can serve as a model to investigate MFS pathology. These findings establish a human iPSC platform for MFS endothelial research and suggest impaired inflammatory resolution as a novel therapeutic target.

## Introduction

Marfan syndrome (MFS, OMIM: 154700) is an autosomal dominant connective tissue disorder, characterized by variants in the FBN1 gene which encodes the extracellular matrix (ECM) protein fibrillin-1 (Zeigler et al., 2021). Fibrillin-1 is a glycoprotein and a main component of ECM microfibrils (Hayward and Brock, 1997). Variants in the FBN1 gene can lead to a reduction of fibrillin-1 bioavailability or impairment of microfibril formation (Mieremet et al., 2022, Jensen et al., 2021). Besides its structural role, fibrillin-1 regulates cell signaling, proliferation and migration (Mariko et al., 2010). Moreover, the fibrillin ECM network captures growth factors, such as transforming growth factor (TGF-β) family members, to be released upon ECM injury (Spanou and Sengle, 2025). Hence, an altered microfibril network may drive uncontrolled release of such growth factors.

MFS has an estimated prevalence of 2–3 per 10,000 (Judge and Dietz, 2005) and is classified as a systemic disease with skeletal, ocular, and cardiovascular implications (Milewicz et al., 2021). Particularly aneurysms and dissections/ruptures of the aortic root and ascending aorta, represent the most severe and life-threatening complications of MFS (J L Murdoch, 1972). Current clinical management include long-term medical treatment with β-blockers or angiotensin-II receptor blockers and potential surgical interventions to stabilize the aorta (Muino-Mosquera et al., 2024). After aortic root and/or ascending aorta replacement surgery, dissection of the thoracic descending aorta remains a risk, likely due to hemodynamic fluctuations (den Hartog et al., 2015).

The human aorta is a multi-layered tissue. The innermost layer (intima), facing the blood and exposed to flow-induced shear forces, contains a single cell layer of endothelial cells (ECs) positioned on a basement membrane and the internal elastic lamina. Below the intima is the media, consisting of vascular smooth muscle cells (VSMCs) in multiple layers divided by elastic lamellae. The outermost layer, the adventitia, is composed of collagen fibers which serve as a natural external stent that are maintained by fibroblasts, and a microvascular network (vasa vasorum) feeding into the medial layer (Hauger and Hordijk, 2024). It is well established that VSMCs play a central role in MFS pathogenesis, exhibiting multiple forms of dysregulation, including altered TGF-β signaling, phenotypic switching, mitochondrial dysfunction, increased ECM and matrix metalloproteinase expression, enhanced production of pro-inflammatory cytokines and cell death (Perrucci et al., 2017).

Additionally, aortic disease in MFS has been increasingly linked to endothelial dysfunction. Patients showed elevated von Willebrand factor (VWF) and thrombomodulin levels (Dirk G. Wilson, 1999) and an increase in aortic stiffness (Sandor et al., 2003, Wanga et al., 2017). Flow-mediated dilation, a noninvasive measurement of endothelial nitric oxide synthase (eNOS) function, was evaluated in patients with MFS, and impaired flow-mediated dilation correlated with enhanced aortic diameters (Takata et al., 2014). Animal models of MFS further showed that EC alignment in the direction of blood flow, junctional organization and EC eNOS expression is impaired in aortae of Fbn1^C1041G/+^ MFS mice (Mieremet et al., 2022). Moreover, a model of aortic microdissection in Fbn1^G234D^ mutant mice showed impaired EC mechanosensing, reduction of flow alignment in the aorta and increased EC-immune cell interaction (Kimura et al., 2025).

Most data on EC dysfunction in MFS are derived either from animal studies, which may lack full translational relevance and require validation in human tissue or cells. To overcome these limitations and to study the role of human MFS ECs in detail, we generated three induced pluripotent stem cell (hiPSC) derived EC lines from MFS patients (iMFS-ECs), each carrying a distinct variant in the *FBN1* gene, as well as 3 healthy control hiPSC derived EC lines (iECs). We find that, compared to control iECs, iMFS-ECs (i) recapitulate MFS phenotypes, such as loss of flow-induced alignment; (ii) show a striking deficiency in recovery from TNF-α-induced loss of barrier function; and (iii) show clear differences in gene expression linked to cytoskeletal and ECM remodeling.

## Results

### Generation of Endothelial Cells from Healthy and MFS hiPSCs

To study the role of EC dysfunction in MFS, we generated MFS patient derived hiPSC-ECs (iMFS-EC; AUMCi016-A, AUMCi017-A, AUMCi018-A), from 3 patients carrying variants in the *FBN1* gene. As controls, we used iECs derived from 3 healthy donors (iECs; SCVI111, SCVI114, GSB-L480) (Figure 1A). Characterization via immunofluorescence staining confirmed that generated iECs show a defined EC-like morphology, depicted by a confluent monolayer with VE-cadherin junctions for iECs (Figure 1B) and iMFS-ECs (Figure 1C). We further quantified the expression levels of the key EC markers VE-cadherin and PECAM-1 and found that ≥98.5% of both iECs and iMFS-ECs were double positive by flow cytometry, indicating highly efficient and homogeneous EC differentiation (Figure 1D). Lastly, we quantified protein expression of VE-cadherin and PECAM-1 via western blot and found that protein expression levels for both markers are comparable between iECs and iMFS-ECs (Figure 1 E, F).

**Figure 1:**
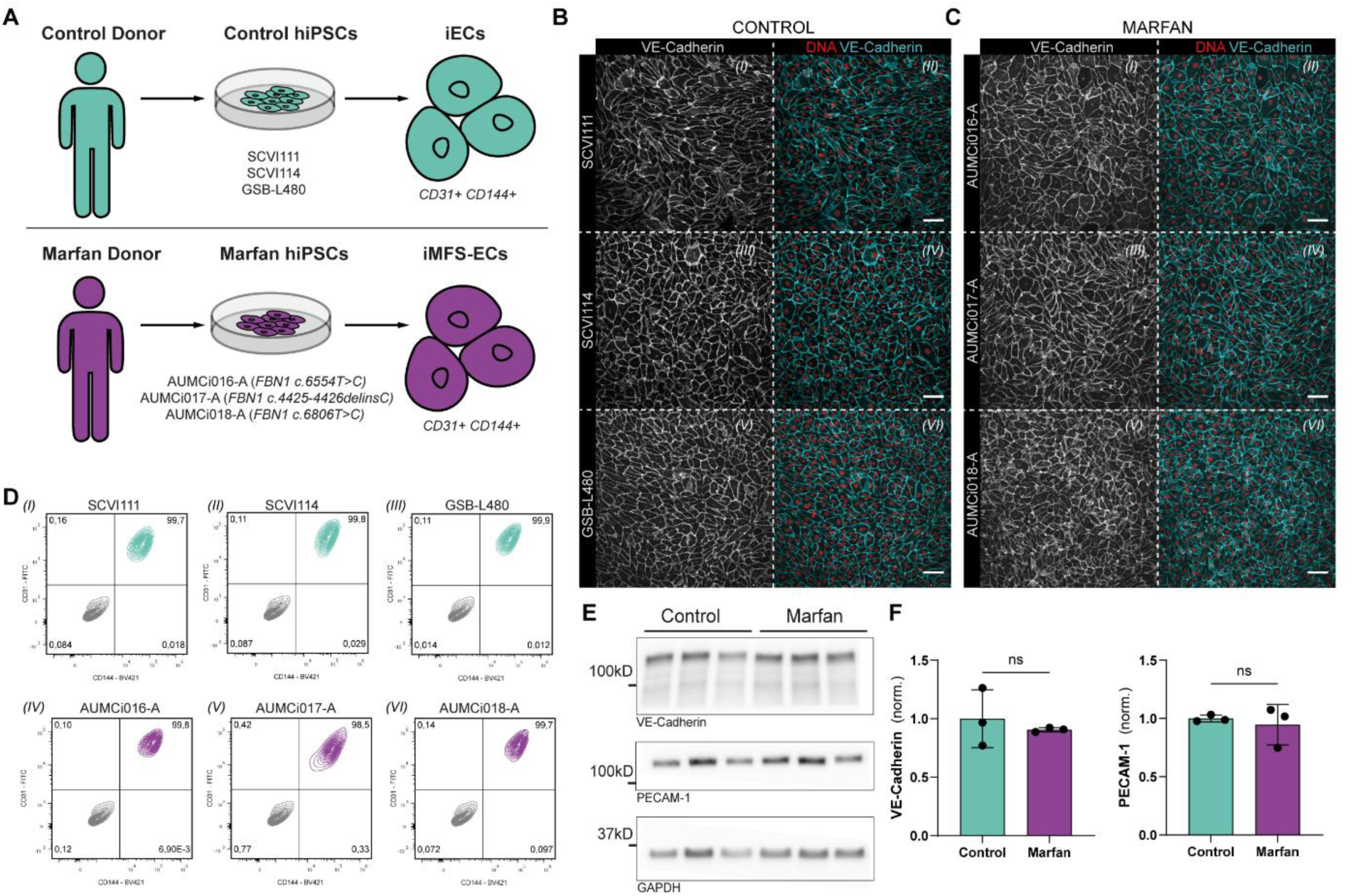
Generation and characterization of hiPSC derived ECs. (A) Schematic overview of the study design. Top: 3 iEC lines (SCVI111, SCVI114, GSB-L480), bottom: 3 iMFS-EC lines (AUMCi016-A, AUMCi017-A, AUMCi018-A) generated in this study. (B) Representative confocal images of iECs immunostained for DNA (red) and VE-Cadherin (cyan). Right panel: VE-Cadherin only, left panel: Overlay of DNA and VE-Cadherin. Scale bar: 100µm. (C) Representative confocal images of iMFS-ECs immunostained for DNA (red) and VE-Cadherin (cyan). Right panel: VE-Cadherin only, left panel: Overlay of DNA and VE-Cadherin. Scale bar: 100µm. (D) Representative bivariate flow cytometry plots showing cells stained with CD31–FITC (y-axis) and CD144-BV421 (x-axis). Quadrants were defined using shared unstained and single-stained controls. Values in each quadrant indicate the percentage of total gated events. (I) – (III) iECs (green), (IV) – (VI) iMFS-ECs (purple). (E) Representative immunoblots showing VE-cadherin (∼120 kDa) expression (top panel), PECAM-1 expression (∼120 kDa) (center panel) and housekeeping gene GAPDH (∼37 kDa) expression (bottom panel) in Control and Marfan samples. Equal amounts of total protein were loaded per lane. (F) Quantification of Western Blot for VE-Cadherin and PECAM-1. Data are normalized against respective GAPDH expression levels and are shown as mean ± SD. Datapoints represent 3 independent iEC lines and 3 independent iMFS-EC lines. Data passed Shapiro–Wilk normality test, groups compared using unpaired two-tailed t-test. ns>0.05.

### iMFS-ECs show reduced flow alignment *in vitro*

Exposure to laminar shear stress (LSS) induces EC elongation and alignment in the direction of flow, which is considered protective (Malek AM, 1999). EC flow responses can be quantified either by changes in cell shape or by LSS induced intracellular polarization. Under flow, the Golgi apparatus is positioned in front of the nucleus, opposite to the direction of flow (Hikita et al., 2018). Previous studies showed that Fbn1^C1041G/+^ and Fbn1^G234D^ mutant mice exhibited reduced aortic EC alignment in the direction of flow (Mieremet et al., 2022). To test whether iMFS-ECs recapitulate these functional aberrations *in vitro*, we cultured iECs and iMFS-ECs in laminar flow chambers and applied LSS of 18 dynes/cm² for 72h (Figure 2A). We then fixed and immunostained the cultures for VE-cadherin, Golgin97 and DNA (Fig. 2B), and analyzed flow alignment using the Polarity–JaM software (Giese et al., 2025).

**Figure 2:**
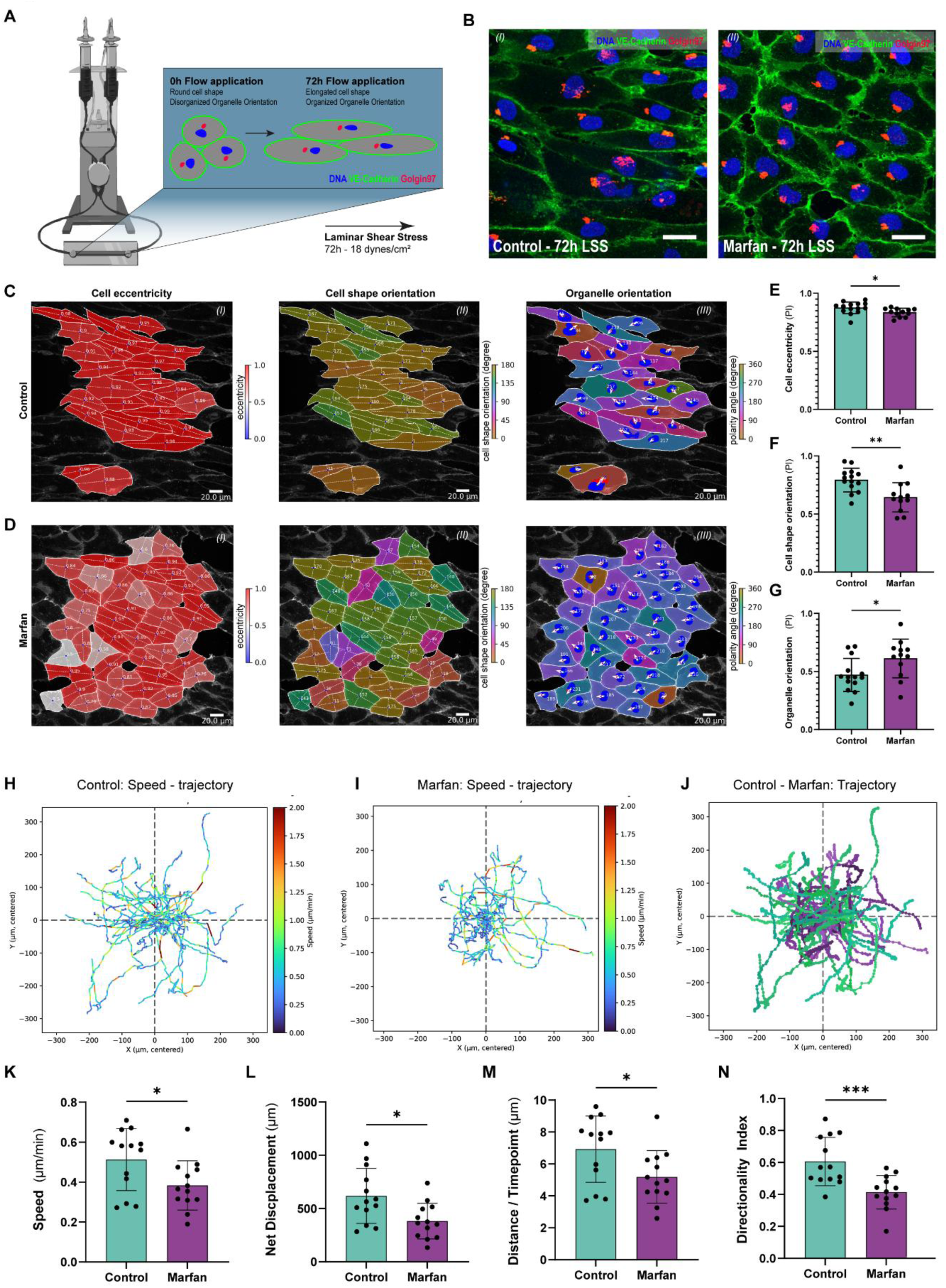
Flow response and migration assessment in hiPSC derived ECs. (A) Schematic overview of experimental set up to study flow response in iECs and iMFS-ECs. Cells were seeded in laminar flow chambers and connected to an ibidi flow unit. LSS was applied for 72h at 18 dynes/cm². (B) Representative confocal images of iECs (I) and iMFS-ECs (II) after exposure to 72h of 18 dynes/cm² LSS (blue: DNA (DAPI), green: VE-Cadherin, red: Golgin-97). Scale bars: 25µm. (C,D) Representative images of Polarity-JaM feature extraction on iECs (C) and iMFS-ECs (D) after LSS exposure. (I) Cell eccentricity, (II) cell shape orientation, (III) organelle orientation. Scale bars: 20µm. Quantification of cell eccentricity (E), cell shape orientation (F), and organelle orientation (G). Datapoints represent the polarity index (PI, 0 – 1) for one image, calculated as average PI per image by the Polarity-JaM Web-app from extracted features generated by the analysis pipeline. 9 independent experiments were performed using two iEC lines (SCVI111, SCVI114) and three iMFS-EC lines (AUMCi016-A, AUMCi017-A, AUMCi018-A). Statistical analyses were performed on pooled individual datapoints per group. Sample numbers: SCVI111 n=8, SCVI114 n=6, AUMCi016-A n=5, AUMCi017-A n=4, AUMCi018-A n=3. Data passed the Shapiro–Wilk normality test and groups were compared using an unpaired two-tailed t-test. Data are shown as mean ± SD. *p<0.05, **p<0.01. (H) Representative trajectory plot of iECs showing cell migration per cell over time (t=12h, 5 min imaging interval). Color code: Speed in µm/min. (I) Representative trajectory plot of iMFS-ECs showing cell migration per cell over time (t=12h, 5 min imaging interval). Color code: Speed in µm/min. (J) Overlayed trajectory plot of iECs and iMFS-ECs showing cell migration per cell over time (t=12h, 5 min imaging interval). Greens: iECs, purples: iMFS-ECs. (K) Quantification of migration speed in µm/min. (L) Quantification of Net Displacement in µm, calculated as the total migrated distance over 12h recording time. (M) Quantification migrated distance per 5 min interval (µm per cell). (N) Quantification of directionality index, defined as net Euclidean distance divided by total migration path length (0–1; 1 = straight-line migration). (K – N): Single cells were analyzed across 7 independent experiments (SCVI111: 43, SCVI114: 36, GSB-L480: 14; AUMCi016-A: 17, AUMCi017-A: 32, AUMCi018-A: 32). Datapoints represent the mean value of one cell line in a single experiment. Statistical analysis was performed on experiment-level means. Data passed Shapiro–Wilk normality test, groups compared using unpaired two-tailed t-test. Data are shown as mean ± SD. ***p<0.005, **p<0.01, *p<0.05.

To confirm overall flow responsiveness, iECs and iMFS-ECs exposed to 72h of LSS were compared with static controls, revealing a clear flow response in both cell types (Supplementary Figure S1). Visual assessment of Polarity–JaM outputs from cells exposed to 72h of LSS indicated that iECs show expected morphological adaptions to flow, such as elongation (Figure 2C-I) and orientation in flow direction (Figure 2C-II), whereas iMFS-ECs depicted a less adapted morphology, showed by reduced elongation (Figure 2D-I) and orientation towards direction of flow (Figure 2D-II). Organelle orientation analysis suggested that both iECs and iMFS-ECs (Fig. 2C,D-III) exhibit a non-random Golgi–nucleus orientation. Quantification of polarity indices confirmed that iMFS-ECs have a reduced cell alignment to flow, depicted by reduced cell eccentricity (Figure 2E) and orientation in the direction of flow (Figure 2F). Interestingly, we found that the organelle orientation in iMFS-ECs was increased over iECs (Figure 2G), suggesting that organelle orientation is not directly coupled to the elongation process.

### iMFS-ECs depict reduced migratory abilities

To determine whether iMFS-ECs display an intrinsic migratory defect, we seeded cells at low density, followed by imaging starting 2h post-seeding at 5-min intervals for 12h. Subsequent cell tracking revealed migratory speed pathways for iECs (Figure 2H) and iMFS-ECs (Figure 2I), and an overlay indicates a decreased migratory ability at single-cell level for iMFS-ECs (Figure 2J). Quantification confirmed this, revealing that iMFS-ECs show reduced migratory speed (Figure 2K), total area migrated over recording time (Figure 2L), migrated distance per imaging timepoint (Figure 2M) and directionality index (Figure 2N). These results confirm the previously reported migratory defect of Marfan iPSC-derived ECs in wound-healing assays (Chen et al., 2025) and show that impaired migration is already evident at the single-cell level in sparsely seeded cultures.

### iMFS-ECs show increased basal endothelial barrier resistance

To further assess the functional integrity of iMFS-ECs *in vitro*, EC barrier function was monitored in real time using electric cell–substrate impedance sensing (ECIS). At equal seeding densities, iMFS-ECs formed a stronger EC barrier than iECs, starting from approximately 5h after seeding and remaining stable for the duration of the recording (Figure 3A). Quantification of resistance values for up to 48h after seeding show that this increased EC barrier in iMFS-ECs is significant between 10h and 48h (Figure 3B).

**Figure 3:**
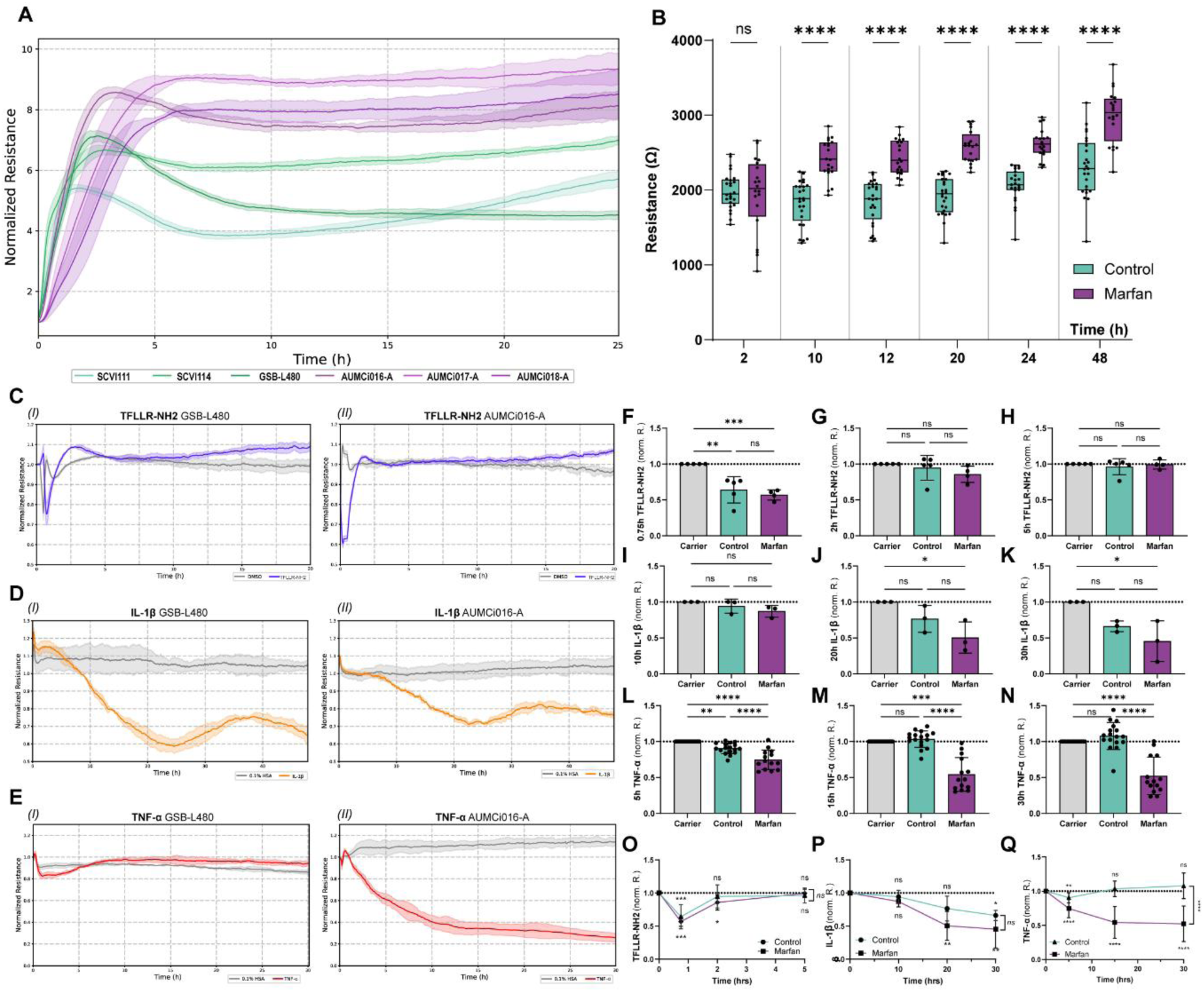
Endothelial barrier evaluation in hiPSC derived ECs. (A) Representative ECIS graph showing endothelial barrier resistance for 25h after cell seeding normalized to t=0h. Greens: iECs, purples: iMFS-ECs. (B) Quantification of endothelial barrier resistance (Ω), for 48h post seeding. Greens: iECs, purples: iMFS-ECs. Resistance values were analyzed across 13 independent experiments. SCVI111 n=13, SCVI114 n=8, GSB-L480 n=4; AUMCi016-A n=7, AUMCi017-A n=10, AUMCi018-A n=3. Statistical analysis was performed on group level means. Data are shown as box plot, whiskers indicate min to max. Data passed Shapiro–Wilk normality test, groups compared using unpaired two-tailed t-test at distinct timepoints. ****p<0.001, ns>0.05. (C–E) Representative ECIS resistance values normalized to the treatment time point following stimulation with TFLLR-NH2 (50µm) (C), IL-1β (100 pg/mL) (D), or TNF-α (10 ng/mL) (E); left: GSB-L480 (I), right: AUMCi016-A (II). (F-H) Quantification of barrier resistance values in iECs (greens) and iMFS-ECs (purples) in response to TFLLR-NH2 (50µM), at 0.75h (F), 2h (G) and 5h (H) post stimulation. Datapoints represent the normalized resistance to a respective vehicle control per cell line and time point. iEC: n=2 (SCVI111), n=2 (GSB-L480), n=1 (SCVI114); iMFS-EC: n=2 (AUMCi017-A), n=1 (AUMCi016-A, AUMCi018-A). (I-K) Quantification of barrier resistance values in iECs (greens) and iMFS-ECs (purples) in response to IL-1β (100 pg/mL) at 10h (I), 20h (J) and 30h (K) post stimulation. Datapoints represent the normalized resistance to a respective vehicle control per cell line and time point. iEC: n=1 (SCVI111, SCVI114, GSB-L480); iMFS-EC: n=1 (AUMCi016-A, AUMCi017-A, AUMCi018-A). (L–N) Quantification of barrier resistance values in iECs (greens) and iMFS-ECs (purples) in response to TNF-α (10ng/mL), at 5h (L), 15h (M) and 30h (N) post stimulation. Datapoints represent the normalized resistance to a respective vehicle control per cell line and timepoint. iEC: n=9 (SCVI111), n=3 (GSB-L480), n=5 (SCVI114); iMFS-EC: n=5 (AUMCi016-A), n=6 (AUMCi017-A), n=3 (AUMCi018-A). (F-N) Statistical analysis was performed on group level mean, data passed Shapiro–Wilk normality test, groups compared using one-way ANOVA. Data are shown as mean ± SD, ****p<0.001, ***p<0.005, **p<0.01, ns>0.05. (R – T) Overview of barrier resistance analysis for TFLLR–NH2 (O), IL-1β (P) and TNF-α (Q) stimulation of iECs (green) and iMFS-ECs (purple) over time. Datapoints represent the averaged normalized resistance to a respective vehicle control for iECs or iMFS-ECs and timepoint, n_iEC-TFLLR-NH2=_5, n_iMFS-EC-TFLLR-NH2=_5, n_iEC-IL-1β=_3, n_iEC-IL-1β=_4, n_iEC-TNF-α=_17, n_iEC-TNF-α=_19. Data are shown as mean ± SD, data passed Shapiro–Wilk normality test, groups compared at each time point against respective vehicle control using one-way ANOVA, with an additional unpaired two-tailed t-test performed between iECs and iMFS-ECs at the final time point (indicated by vertical brackets). ****p<0.001, ***p<0.005, **p<0.01, ns>0.05.

### iMFS-ECs exhibit barrier kinetics comparable to iECs upon PAR1 activation and IL-1β stimulation

To further elucidate the EC barrier properties of iMFS-ECs, we examined their response to inflammatory stimuli, known to transiently disrupt EC barrier integrity. To that end, we treated iMFS-ECs and iECs with TFLLR-NH2 (Thrombin mimicking PAR1 activating peptide), Interleukin-1β (IL1β) or tumor necrosis factor-α (TNF-α), 72h after seeding and formation of a stable barrier. Upon treatment with 50µM TFLLR-NH2, we observed that iMFS-ECs and iECs show a transient drop in barrier integrity that is restored ∼2.5h post-treatment (Figure 3C-I, 3C-II). Quantification further confirms a significantly decreased barrier integrity for both iMFS-ECs and iECs at 0.75h (Figure 3F), and a restoration of the barrier integrity 2h post-treatment (Figure 3G), which remains stable for at least 5h (Figure 3H). At all timepoints, there was no significant difference in response to TFLLR-NH2 between iMFS-ECs and iECs (Figure 3O). Upon 100pg/mL IL-1β stimulation, both iECs and iMFS-ECs show a loss in barrier integrity that becomes evident after ∼5h and is partly restored after ∼30h (Figure 3D-I, 3D-II). Quantification of barrier kinetics shows that the barrier remains stable for the first 10h post IL-1β stimulation (Figure 3I), is decreased after 20h (significant for iMFS-ECs) (Figure 3J) and is significantly reduced for both iECs and iMFS-ECs after 30h (Figure 3K). Similar to TFLLR-NH2, we did not observe any significant difference in response to 100pg/mL IL-1β between iECs and iMFS-ECs (Figure 3P).

### iMFS-ECs show sustained barrier loss upon TNF-α stimulation

Lastly, we analyzed the response to 10ng/mL TNF-α stimulation. We found that iECs have a transient loss of barrier integrity at ∼1h, that was restored ∼5h post TNF-α stimulation. For iMFS-ECs, however, we observed that stimulation with TNF-α resulted in a strong reduction of the EC barrier, with an onset at ∼1h post-treatment and a sustained loss in EC barrier integrity for at least 30h post-treatment (Figure 3E-I, 3E-II). Quantification of barrier kinetics confirmed that both iMFS-ECs and iECs respond to 10ng/mL TNF-α with a significant drop in barrier integrity 5h post treatment (Figure 3L). This drop is restored for iECs 15h post treatment, but not for iMFS-ECs (Figure 3M). 30h after stimulation with TNF-α, iECs depict a stable EC barrier, whereas the barrier remains significantly impaired for iMFS-ECs (Figure 3N,Q). To explore whether the sustained TNF-α-induced loss of barrier integrity in iMFS-ECs was accompanied by altered inflammatory activation, iECs and iMFS-ECs were exposed to TNF-α (10 ng/mL) for 5 h or 24 h and assessed by confocal immunocytochemistry. Exploratory evaluation of VE-cadherin, ICAM-1, and F-actin suggested TNF-α-associated changes in both cell types, including apparent cell elongation, ICAM-1 upregulation, and stress-fiber formation. The imaging did not reveal an obvious iMFS-EC-specific difference in these markers under the conditions examined (Supplementary Fig. S2).

### Transcriptomic comparison of iECs and iMFS-ECs

To further investigate the gene expression differences in iECs and iMFS-ECs in response to TNF-α, we performed bulk RNA sequencing. For each EC line (3 iEC lines, 3 iMFS-EC lines), we sequenced a sample 24h after stimulation with TNF-α and a vehicle control. Bulk RNA sequencing profiled transcriptional changes across approximately 20000 expressed genes per sample. To verify EC identity at the transcriptional level, we found that key EC genes were highly expressed (>10,000 reads) at comparable levels in iECs and iMFS-ECs under vehicle control conditions, including PECAM-1, CDH5, MCAM, ESAM, ENG, and VWF (Figure 4A). Notably, we found several junction-associated genes upregulated in iMFS-ECs under basal conditions, including CLDN5, CGN, CDH4 (significant), and CLDN11 (significant) (Figure 4D), further strengthening the hypothesis that increased junction stability results in the increased EC barrier function of iMFS-ECs at baseline (Figure 3H). TJP1 was found to be upregulated in iECs. In total, 331 genes were found to be significantly differentially expressed between iECs and iMFS-ECs under vehicle control conditions and a similar number (262 genes) following TNF-α stimulation. TNF-α elicited a pronounced transcriptional response within both genotypes, leading to significantly differential expression of 2006 genes in iECs (Figure 4C) and 1831 genes in iMFS-ECs (Figure 4D) when comparing vehicle control and TNF-α conditions. Consistent with these findings, principal component analysis showed that TNF-α treatment accounted for the dominant source of variance along Dim1 (29,3%), while genotype contributed more modestly to separation along Dim2 (14,9%) (Supplementary Figure S3).

**Figure 4:**
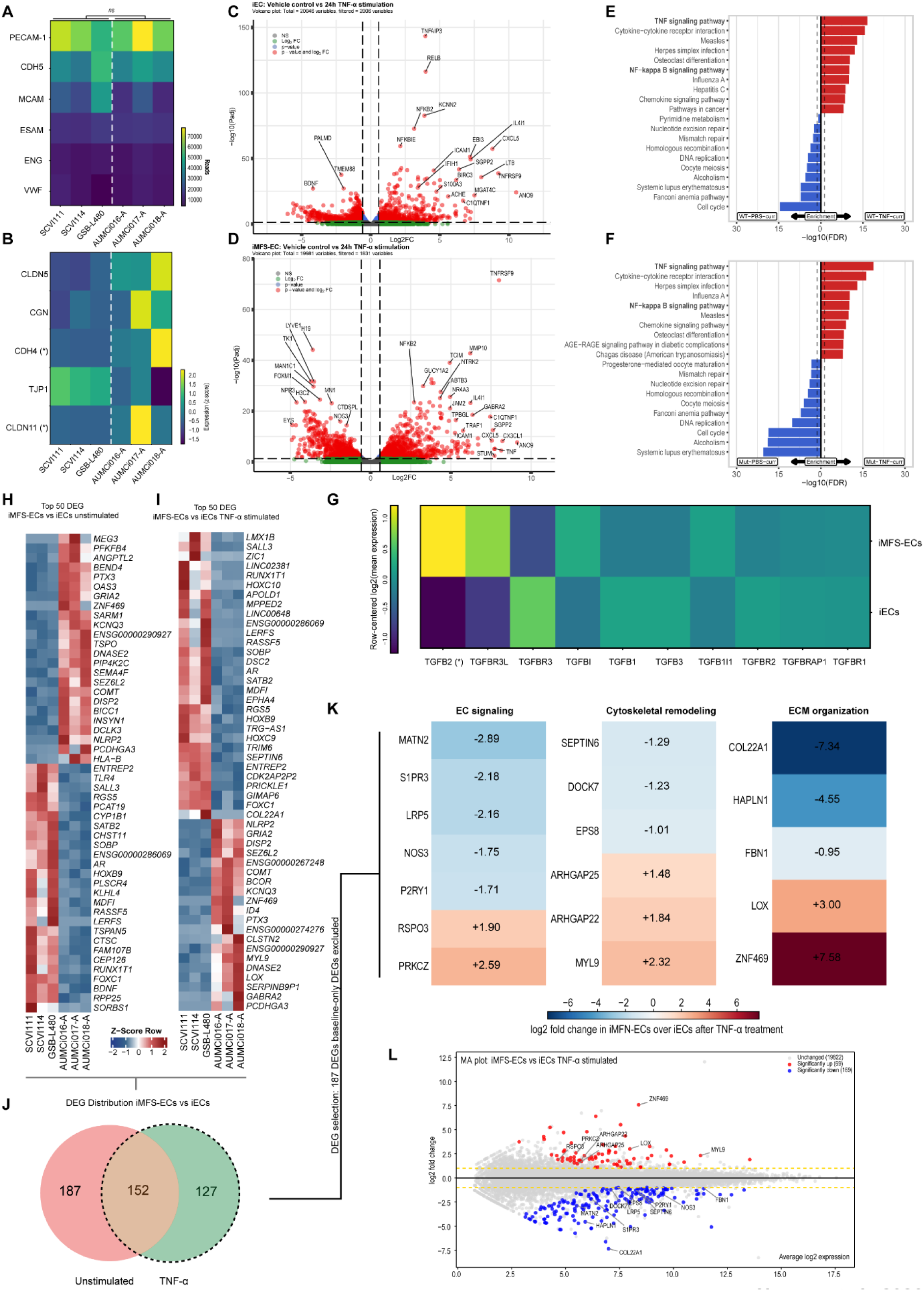
Transcriptomic profiling of iECs and iMFS-ECs after 24 h TNF-α stimulation. (A) Heatmap visualizing gene expression plotting total reads of key EC genes for iECs_Vehicle-Control_ and iMFS-ECs_Vehicle-Control_. padj>0.05 for all genes. (B) Heatmap visualizing gene expression of z-scored reads of junctional genes CLDN5, CGN, CDH4, TJP1 and CLDN11 for iECs_Vehicle-Control_ and iMFS-ECs_Vehicle-Control_. (*)=padj <0.05 (C) Volcano plot showing differential gene expression comparing iECs_Vehicle-Control_ and iEC_TNF-α_. Total recorded variables: 20046, significantly differently expressed: 2006. p-value threshold: p<0.05 (blue), fold change (FC) threshold: |log2(FC)|>0.585 (green). Significant genes above p-value and FC threshold are visualized as red dots. (D) Volcano plot showing differential gene expression for iMFS-ECs, comparing comparing iMFS-ECs_Vehicle-Control_ and iMFS-EC_TNF-α_. Total recorded variables: 19981, significantly differently expressed: 1831. p-value threshold: p<0.05 (blue), FC threshold: |log2(FC)|>0.585 (green). Significant genes above p-value and FC threshold are visualized as red dots. (E) Direction-specific KEGG pathway enrichment comparing iECs_Vehicle-Control_ and iEC_TNF-α_. Upregulated (red) and downregulated (blue) DEGs were analyzed separately using KOBAS with the KEGG PATHWAY database. The dashed line indicates significant enrichment (FDR-adjusted *p*<0.05). (F) Direction-specific KEGG pathway enrichment comparing iMFS-ECs_Vehicle-Control_ and iMFS-EC_TNF-α_. Upregulated (red) and downregulated (blue) DEGs were analyzed separately using KOBAS with the KEGG PATHWAY database. The dashed line indicates significant enrichment (FDR-adjusted *p*<0.05). (G) Row-centered log2 expression heatmap visualizing gene expression of genes in the TGFβ pathway for iECs_Vehicle-Control_ and iMFS-ECs_Vehicle-Control_. (*)=padj<0.05. (H) Row-wise z-scored heatmap showing the top 50 most significantly differentially expressed genes between iEC_Vehicle-control_ and iMFS-EC_Vehicle-control_ ranked by adjusted p-value (padj, smallest to largest) (I) Row-wise z-scored heatmap showing the top 50 most significantly differentially expressed genes between iEC_TNF-α_ and iMFS-EC_TNF-α_ ranked by adjusted p-value (padj, smallest to largest) (J) Venn diagram that includes DEGs of iEC_Vehicle-Control_ versus iMFS-EC_Vehicle-Control_ comparison, and DEGs of iEC_TNF-α_ versus iMFS-EC_TNF-α_ comparison. Left (pink): DEGs in iEC_Vehicle-Control_ versus iMFS-EC_Vehicle-Control_ comparison (187). Middle (orange): DEGs both in iEC_Vehicle-Control_ versus iMFS-EC_Vehicle-Control_ comparison, and iEC_TNF-α_ versus iMFS-EC_TNF-α_ comparison (152). Right (green): DEGs in iEC_TNF-α_ versus iMFS-EC_TNF-α_ comparison (127). (K) Heatmap of selected genes for endothelial signaling, cytoskeletal remodeling, and ECM organization relevant genes in iMFS-EC_TNF-α_. Genes were grouped by literature relevance and ordered within each group from lowest to highest log2 FC. Red: upregulation, blue: downregulation (relative to iEC_TNF-α_). (L) MA plot showing gene selection shown in (K) for iEC_TNF-α_ versus iMFS-EC_TNF-α_. X-axis: Average log2 expression, y-axis: log2 FC. Genes are classified as upregulated (blue), downregulated (red), or unchanged (gray), based on a log2 FC cutoff of ± 0.5 and a p-value threshold of 0.05. Dashed lines: ± 0.5 log2 FC thresholds, yellow lines: log2 FC=0.

### TNF-α upregulates NF-kB and TNF-α signaling pathways in iMFS-ECs and iECs

To identify biological processes associated with TNF-α stimulation in iECs, we performed KEGG pathway enrichment analysis. Gene set enrichment analysis (GSEA) revealed significant enrichment of multiple pathways among both up- and downregulated genes (FDR < 0.5). Amongst the pathways upregulated after TNF-α stimulation were the TNF-α- and NF-kB signaling pathways (Figure 4E). We then performed the same analysis on iMFS-ECs, comparing TNF-α stimulation with vehicle control gene expression, and found a similar pattern in the GSEA, including TNF-α- and NF-kB pathway upregulation (Figure 4F). This analysis supports our previous findings showing that the canonical response to TNF-α stimulation in iMFS-ECs is comparable to iECs. Moreover, expression of ICAM1 and NF-kB pathway genes (NFKB2, NFKBIA, NFKBIB, NFKB1, NFKBIZ, NFKBID, NFKBIL1, NFKBI) is not significantly different between iECs and iMFS-ECs after stimulation with TNF-α (Supplementary Figure S4).

### TGFB2 is significantly upregulated in iMFS-ECs

Mutations in the *FBN1* gene alter TGF-β bioavailability and downstream signaling. We thus investigated the TGF-β family members. In iMFS-ECs, TGFB2 is significantly upregulated in comparison to iECs, whereas the expression of TGFBR3, TGFB1I1, TGFB1, TGFBR2, TGFBR1, TGFB3, TGFBI, TGFBRAP1 and TGFBR3L was comparable (Figure 4G). We furthermore investigated whether the downstream SMAD pathway was dysregulated in iMFS-ECs and found that inhibitory SMAD6 was significantly downregulated in iMFS-ECs. SMAD1-5, 7 and 9 were not significantly different between iECs and iMFS-ECs, however showed a trend towards decreased SMAD2,3,5,7 and 9 gene expression (Supplementary figure S5).

### Differentially expressed genes between iMFS-ECs and iECs following TNF-α stimulation

We next distinguished genes that are specifically associated with TNF-α stimulation from those already differentially expressed between iECs and iMFS-ECs under basal conditions. To this end, we compared genes differentially expressed between iECs and iMFS-ECs under vehicle control conditions (Figure 4H) with those differentially expressed following TNF-α stimulation (Figure 4I). Figure 4H,I show the top 50 DEGs. Venn diagram analysis on the total number of DEGs identified 187 genes that were differentially expressed exclusively under basal conditions and were therefore excluded from subsequent analyses. In contrast, 152 genes were differentially expressed under both basal and TNF-α-stimulated conditions, while 127 genes became differentially expressed only upon TNF-α stimulation (Figure 4J). Subsequent analyses focused on the combined set of genes that were differentially expressed following TNF-α stimulation, including both the TNF-α-specific genes and those overlapping between basal and stimulated conditions.

### Functional Classification of Endothelial Barrier-Associated DEGs

We then performed a targeted, literature-guided annotation of the selected DEGs. As we previously established that the initial inflammatory response following TNF-α stimulation was comparable between iECs and iMFS-ECs, we aimed to identify alternative mechanisms that could contribute to EC barrier loss. Based on established determinants of EC barrier integrity, we established 3 categories: EC signaling, cytoskeletal remodeling, and ECM organization (Figure 4K). Genes from the predefined DEG set were subsequently evaluated for reported roles in EC biology and barrier regulation, and those with clear literature-supported relevance were assigned to these categories, while genes without an evident connection to these processes were not further classified.

To the EC signaling group, we assigned MATN2, S1PR3, LRP5, NOS3, P2RY1, R-RSPO3 and PRKCZ. In the cytoskeletal remodeling group, we assigned SEPTIN6, DOCK7, EPS8, ARHGAP25, ARHGAP22 and MYL9. Finally, we assigned to the ECM organization group COL22A1, HAPLN1, FBN1, LOX and ZNF469 (Figure 4J). An MA plot was generated to visualize the relationship between differential expression and average transcript abundance, demonstrating that the identified candidate genes were robustly expressed across samples, with average log2 expression levels above 4.5 (Figure 4L).

### Functional Interpretation of Barrier-Associated DEGs

We next analyzed the differential expression profiles of genes within each functional group to assess the directionality of transcriptional changes in EC barrier-associated pathways. Within the EC signaling group, we observed that several identified genes linked to PI3K/AKT signaling pathways were downregulated in iMFS-ECs, including MATN2 (Liu et al., 2024), S1PR3 (Wilkerson and Argraves, 2014), P2RY1 (Cabou and Martinez, 2022) and NOS3 (also known as eNOS) (Peng et al., 2010), and PI3K/AKT signaling is known to promote endothelial barrier integrity (Huang et al., 2016b, Gunduz et al., 2019). Moreover, the Wnt receptor LRP5 was found to be downregulated in iMFS-ECs, which has been previously linked to impaired vascular development in the retina (Huang et al., 2016a). Amongst the genes we classified in the EC signaling group, RSPO3 and PRKCZ were found to be upregulated in iMFS-ECs. RSPO3 is a Wnt pathway ligand, and was previously linked to EC barrier disruption *in vitro* (Skaria et al., 2018), and EC recovery after sepsis (Zhang et al., 2024). PRKCZ, encoding for PKCζ, is a kinase that was shown to promote apoptosis in ECs through JNK-caspase 3 mediated signaling (Kim et al., 2012) and its inhibition was found to promote EC barrier integrity during inflammation (Lin et al., 2018).

Of the genes we associated with cytoskeletal remodeling, we found a majority linked to Rac1 regulation, a Rho GTPase that stabilizes the EC barrier upon activation (Waschke et al., 2006). Rho GTPases are molecular switches that regulate cytoskeletal dynamics, cell polarity, migration, and EC barrier function, with RhoGEFs activating them by promoting GDP-to-GTP exchange and RhoGAPs inactivating them by accelerating GTP hydrolysis (Mosaddeghzadeh and Ahmadian, 2021). Within our dataset, we found the Rac1 RhoGEF DOCK7 (Kukimoto-Niino et al., 2019) downregulated and the Rac1 RhoGAPs AHRGAP22 (Mori et al., 2014) and AHRGAP25 (Huang et al., 2021) upregulated in iMFS-ECs, suggesting decreased levels of active Rac1 in iMFS-ECs upon TNF-α stimulation. SEPTIN6 is a member of the Septin family, GTP-binding proteins that form cytoskeletal like structures (Mostowy and Cossart, 2012). Septins interact closely with Rho GTPase signaling pathways and act as scaffolding elements that coordinate actin and microtubule organization, thereby contributing to cytoskeletal stability and EC junction integrity (Dolat et al., 2014). SEPTIN6, which is linked to Cdc42, a Rho GTPase with barrier-promoting effects similar to Rac1, was found to be downregulated in iMFS-ECs following TNF-α stimulation (Gérard Joberty, 2001). We furthermore found elevated levels of MYL9 in iMFS-ECs, which is associated with RhoA/ROCK-mediated actomyosin contraction (Wojciak-Stothard and Ridley, 2002) and implicated in age-related vascular barrier impairment (Shehadeh et al., 2011). Lastly, we found reduced levels of the actin dynamics regulator EPS8, which was linked to increased vascular leakage *in vivo* (Giampietro et al., 2015).

Within the ECM organization category, we found COL22A1, HAPLN1 and FBN1 downregulated. COL22A1 has been shown to play a role in vascular maintenance and its loss was correlated to aneurysm formation (Ton et al., 2018). HAPLN1 serves as link between hyaluronic acid and proteoglycans in the ECM. Age related decreased HAPLN1 levels were linked to EC senescence (Zhou et al., 2023) and impaired vascular integrity in melanoma (Zhou et al., 2023). Interestingly, reduced FBN1 expression in TNF-α-stimulated iMFS-ECs was found, consistent with the underlying loss-of-function FBN1 mutations in our patient-derived cell lines. It is furthermore established that fibrillin-1 is crucial for cellular signaling and fibrillin-1 impairment contributes to blood-brain barrier breakdown in mice (Van der Donckt et al., 2015). Within this group, LOX and ZNF469 were upregulated in iMFS-ECs upon TNF-α stimulation. LOX is an aneurysm related gene, since it is essential in crosslinking collagens and elastin to form an integrated ECM. As such it plays a role in tumor angiogenesis and EC migration (Baker et al., 2013). ZNF469 was shown to regulate collagen gene expression in human hepatic stellate cells (Steinhauser et al., 2025) and dermal fibroblasts (Charoenthanakitkul et al., 2025). ZNF469 mutations are linked to connective tissue disorders (Steinle et al., 2022), suggesting that its upregulation in iMFS-ECs may reflect enhanced ECM remodeling that could contribute to EC barrier dysfunction.

In conclusion, the gene set analysis provides a mechanistic framework, suggesting that iMFS-ECs respond canonically to TNF-α mediated inflammation, however 24h after the insult, barrier maintaining pathways like the PI3K/AKT and Wnt pathway remain dysregulated. The cells remain in a contractile state, possibly further amplified by impaired Rac1/enhanced RhoA signaling. Additionally, the enrichment of matrix remodeling-associated gene sets suggests an aberrant ECM regulatory state. Collectively, these processes may contribute to the impaired barrier phenotype observed in iMFS-EC 24h post TNF-α stimulation.

### Rho-kinase inhibition partly rescues TNF-α induced barrier loss in iMFS-ECs

To follow up on the hypothesis that iMFS-ECs respond to TNF-α with increased contraction and thus a reduced restoration of the EC barrier, we aimed to rescue this phenotype. First, we investigated if the Wnt-pathway is involved. We analyzed whether Wnt pathway inhibition could improve barrier integrity by pretreating iMFS-ECs with the Wnt inhibitor C59 (2µM) for 2h prior to TNF-α stimulation. However, we could not rescue the TNF-α-induced barrier loss (Supplementary figure S6). Then, we tested for a disturbed Rho kinase–mediated cytoskeletal regulation, which is tightly balanced; while Rac1 promotes cell spreading, RhoA/B activate Rho-kinase (ROCK) to induce cell contraction. To test whether Rac1 activation could rescue the observed phenotype, Epac/Rap1 signaling was induced to promote EC barrier stabilization in a Rac1 dependent manner and counteracting actomyosin-driven contraction. To that end, we pretreated iMFS-ECs with 2µM 8-pCPT-2’-O-Me-cAMP (“007cAMP”) for 1h. We observed that in both iECs and iMFS-ECs, pretreatment with 007cAMP did not impact the barrier (Figure 5C, D). Specifically, in iMFS-ECs we did not see a rescue of TNF-α-induced barrier loss. Since TNF-α is known to increase reactive oxygen species (ROS) in ECs in a Rac-dependent manner (Marcos-Ramiro et al., 2014), we investigated whether iMFS-ECs exhibit increased ROS production upon TNF-α stimulation. ROS levels were modestly elevated compared to carrier-treated controls 7h after treatment, an effect that was not observed in iECs. However, after 24h ROS levels had returned to baseline (Supplementary Figure S7). Combinatorial treatment with the ROS scavenger N-Acetylcysteine and TNF-α could not rescue the sustained barrier loss in iMFS-ECs (Supplementary Figure S8).

**Figure 5:**
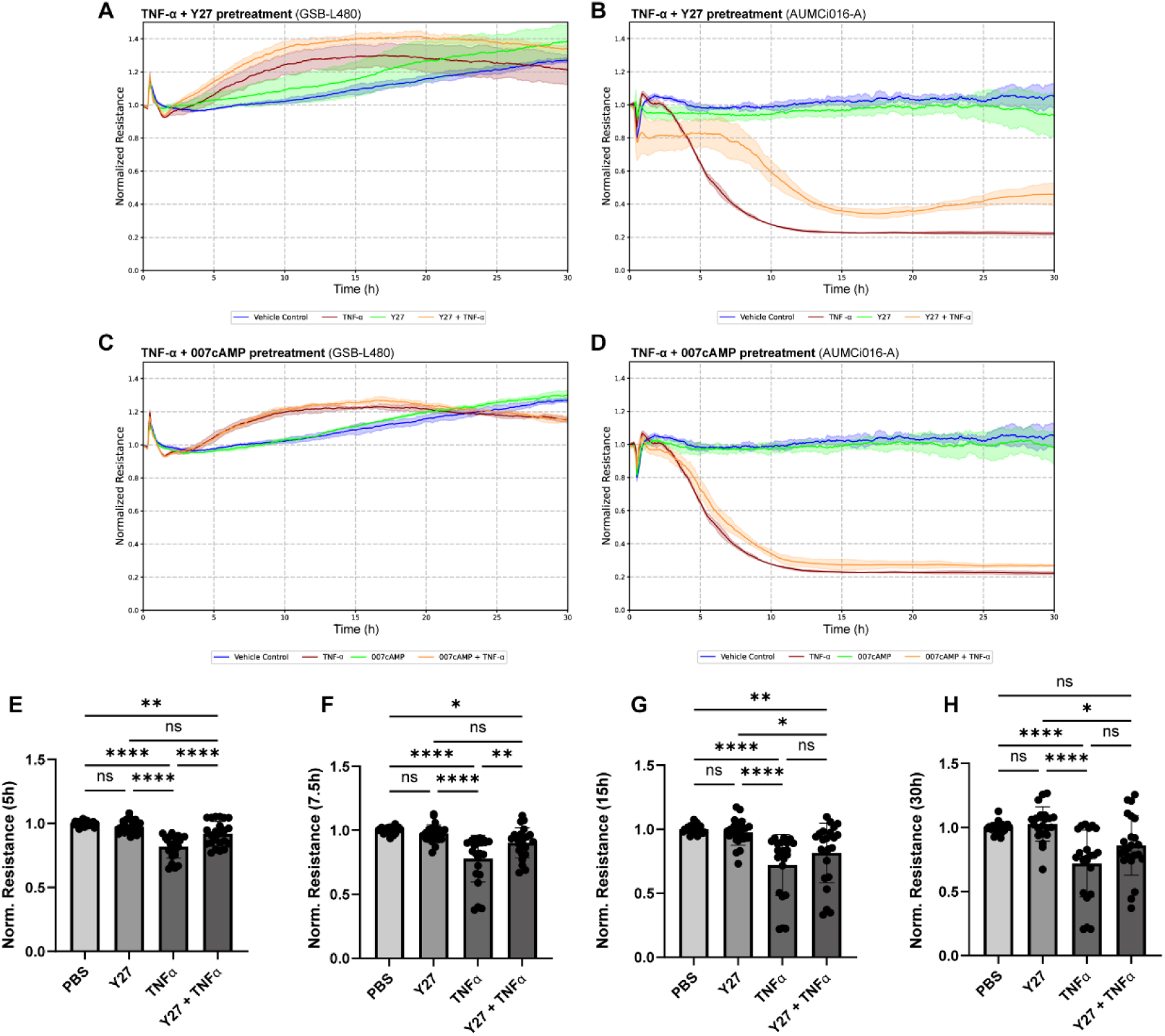
Partial restoration of endothelial barrier function by pharmacological intervention. (A-D) Representative ECIS resistance values normalized to the treatment time point following stimulation with 10ng/mL TNF-α and 1h pretreatment with 10µM Y-27632 (A: iECs, B: iMFS-ECs); or 1h pretreatment with 2µM 8-pCPT-2’-O-Me-cAMP (C: iECs, D: iMFS-ECs). Blue: vehicle control, green: pretreatment only, red: TNF-α, orange: pretreatment and TNF-α. (E-G) Quantification of barrier resistance values in iMFS-ECs (AUMCi016-A) after treatment with TNF-α, or pretreatment with Y-27632 or 8-pCPT-2’-O-Me-cAMP for 1h following TNF-α treatment. (E) 5h, (F) 7.5h, (G) 15h and (H) 30h post TNF-α treatment. Data points represent the normalized endothelial resistance measured in technical triplicates from three independent iMFS-EC lines (AUMCi016-1, n=1; AUMCi017-A, n=3; AUMCi018-A, n=3), with values normalized to the respective carrier control at each time point. Statistical analysis was performed on the group-level means. Data passed the Shapiro–Wilk normality test and groups were compared using one-way ANOVA. Data are presented as mean ± SD. \*\*\*\**p* < 0.001, \*\**p* < 0.005, \**p* < 0.01, *p* < 0.05, ns, not significant (*p* > 0.05).

Then, we investigated if inhibition of RhoA-mediated signaling could attenuate the TNF-α-induced contractile response. The iMFS-ECs were pretreated for 1h with 10µM ROCK inhibitor Y-27632 (“Y27”) prior to stimulation with 10ng/mL TNF-α. Now, we observed that pretreatment with Y27 ameliorates the TNF-α-induced barrier drop in iMFS-ECs, whereas treatment with Y27 did not have a clear effect on unstimulated iMFS-ECs (Figure 5B). In iECs, Y27 did not affect the barrier in the presence or absence of TNF-α (Figure 5A). We quantified these observations for iMFS-ECs at timepoints 5h (Figure 5E), 7.5h (Figure 5F), 15h (Figure 5G) and 30h (Figure 5H) post TNF-α treatment in iMFS-ECs, and found that blockade of RhoA-induced contraction significantly rescued the TNF-α-induced drop in EC barrier in iMFS-ECs at timepoints 5h and 7.5h.

## Discussion

In the present study, we explored the phenotype of hiPSC-derived ECs from 3 Marfan patients, each harboring a different mutation in the FBN1 gene. Each hiPSC line could be differentiated into ECs. The iMFS-ECs were morphologically similar to healthy control hiPSC-derived iECs, and expressed EC markers CD31 and CD144 comparably. Functionally, iMFS-ECs retained key characteristics of EC behavior, including the ability to form confluent monolayers *in vitro,* accompanied by a high and stable EC barrier resistance. We then aimed to recapitulate previously reported MFS EC phenotypes observed in murine Marfan syndrome models, using our human hiPSC-EC *in vitro* system.

Murine Marfan syndrome models demonstrated impaired EC cell alignment and elongation in the direction of flow by *en face* staining of the aortic endothelium (Mieremet et al., 2022), which could be mimicked by the iMFS-ECs *in vitro.* However, the golgi – to – nuclei angles were not reduced by flow, but rather enhanced in iMFS-ECs in comparison to iECs. It has previously been shown that ECs respond to shear stress in a magnitude-depending manner (Sonmez et al., 2020). We hypothesize that iMFS-ECs, due to the lack of alignment in the direction of flow, experience a net higher shear stress magnitude in comparison to iECs when exposed to the same dyne/cm² shear input, due to their different cell shape. Assuming the mechanosensing response is not defective, this suggests that changing cell shape may be compromised in iMFS-ECs. This is in line with previous reports (Chen et al., 2025) and our results on decreased cell migration of iMFS-ECs, where changing shape is essential. Moreover, upon TNF-α-induced inflammatory stimulation, iMFS-ECs do not recover from the EC barrier loss, while these cells respond normally to IL-1β and PAR1 agonist TFLLR-NH2. This selective differential response of the MFS cells was associated with dysregulated PI3K/AKT signaling, ECM remodeling, and impaired cytoskeletal dynamics, where the latter is related to shape changes and will be discussed later. iMFS-ECs showed higher baseline barrier resistance and transcript levels of Claudin-5, CGN, CDH4, and CLDN11. We conclude that increased tight and adherens junction levels may partly underlie this increased baseline barrier. This is further supported by single-cell RNA-sequencing data from human Marfan aortic tissue showing elevated tight and adherens junction gene expression in a subset of endothelial cells *in vivo* (Dawson et al., 2021), raising the possibility that iMFS-ECs adopt a remodeling-associated endothelial phenotype.

On a transcriptomic level, iMFS-ECs show high similarity to iECs with intact canonical TNF-α-responsive pathways, including ICAM1 upregulation and NF-κB activation. An increased inflammatory phenotype of the MFS aorta is linked to disturbed TGF-β signaling rather than increased levels of other cytokines such as TNF-α (Radonic et al., 2012). In our dataset, we found an upregulation of TGFB2 and downregulation of SMAD6 gene expression in iMFS-ECs. TGFB2 encodes for TGF-β2, which is important for the development of the aorta and outflow tract (L. Philip Sanford, 1997). Reduced TGFB2 expression in a MFS background enhanced aortic aneurysm severity in mice (Deleeuw et al., 2023), showing that TGFB2 is essential for aorta homeostasis. SMAD6 is an inhibitor of the bone morphogenetic protein (BMP) pathway, which is part of the TGF-β family signaling cascades. Since the different TGF-β family growth factors and receptors are interconnected, altered signaling may impact cellular state. It is well established that fibrillin-1 supports the sequestration and regulation of latent TGF-β complexes, and pathogenic *FBN1* variants have been associated with changes in TGF-β signaling (Sengle and Sakai, 2015). Yet, we did not observe broad transcriptional activation of canonical TGF-β target genes. This may be explained by the fact that canonical TGF-β/SMAD signaling is extensively regulated at the post-translational level (Massague, 2012).

TNF-α is known to impair EC barrier integrity through cytoskeletal remodeling and junctional destabilization. The sustained phenotype observed in iMFS-ECs suggests that these cells have impaired pathways to resolve barrier disruption. This interpretation is further supported by the upregulation of MYL9, a regulator of myosin-mediated contractility, together with transcriptional changes indicative of suppressed Rac1 signaling, including downregulation of Rac1 activators and upregulation of Rac1 deactivators 24h after stimulation with TNF-α. As Rac1 activity is essential for maintaining cortical actin organization and endothelial junction stability, reduced Rac1 signaling may shift the cytoskeletal balance toward a RhoA/ROCK-dominant contractile state that promotes stress fiber formation and junctional disruption. Indeed, we showed that ROCK inhibition partially rescued the TNF-α-induced loss of endothelial barrier integrity in iMFS-ECs, as evidenced by a reduced decline in barrier function up to at least 7.5 h following TNF-α treatment. However, this protective effect was no longer observed at later time points (15–30 h post-treatment). The incomplete rescue of the phenotype suggests that additional dysregulated mechanisms contribute to the barrier defect. Furthermore, it is possible that the inhibitory activity of the compound diminishes over prolonged treatment, thereby limiting its efficacy at later time points. Interference studies showed that Rap1-signaling, the Wnt pathway or ROS production were no major drivers in the observed TNF-α-induced barrier disruption in iMFS-ECs. We did not detect major differences in canonical TNF-α-responsive inflammatory signaling pathways at either the protein or transcriptomic level, which suggests that the observed EC dysfunction may primarily arise from altered cytoskeletal and mechanotransductive responses rather than exaggerated classical inflammatory activation. In addition, the altered ECM mechanics resulting from fibrillin-1 fiber network dysfunction may enhance mechanosensitive activation of latent TGF-β complexes, thereby contributing to disturbed TGF-β signaling in MFS, establishing a positive feedback loop between matrix dysregulation, cytoskeletal tension, and EC dysfunction.

In summary, we show that hiPSCs carrying a *FBN1* mutation serve as valid model to study mechanisms underlying MFS *in vitro*. These iMFS-ECs recapitulate key EC features such as a stable barrier and marker expression, and mimic disease phenotypes such as alignment with flow as observed *in vivo*. We identified that iMFS-ECs display a sustained loss of barrier integrity after stimulation with TNF-α, which was associated with changes in the PI3K/AKT and Wnt pathways, cytoskeletal regulation, ECM remodeling, and could be partially rescued by ROCK inhibition. These data highlight a critical role for ECs in aortic disease in MFS and identify the vascular endothelium as potential target for future therapeutic strategies.

## Resource availability

### Lead contact

Requests for additional information, resources, or reagents should be directed to the corresponding authors P. C. Hauger, V. de Waard. or P.Hordijk.

### Materials availability

This study did not generate new unique reagents. Requests for additional information or materials should be addressed to the lead contacts.

## Data and code availability

Raw data for RNA Sequencing experiments at NCBI GEO: https://www.ncbi.nlm.nih.gov/geo/query/acc.cgi?acc=GSE335638 *reviewer access token: oxqfyscyrtwpdgt*

Custom code used for analysis of cell trajectories is provided as Python script file in the Supplemental Information accompanying this submission.

## Acknowledgments

The authors thank Prof. Joseph C. Wu from Stanford University for providing the hiPSC line SCVI-111, SCVI114 and GSB-L480. We thank Mike de Kok from Amsterdam UMC for support and discussions regarding data handling and analysis of RNA sequencing data. We furthermore thank Dr. Giulia Bergamaschi from the MCCF Microscopy and Cytometry Core Facility at Amsterdam UMC for support with microscopy data acquisition and data analysis.

## Authors contribution

PCH conceived the study, performed experiments, analyzed data, and wrote the manuscript. GD, LS, CK, and MCO contributed to data acquisition and analysis. JWB, VdW, and PLH provided supervision and contributed to study design and interpretation of results. VdW and PLH supervised the project. All authors reviewed and approved the final manuscript.

---

**Table S1:**
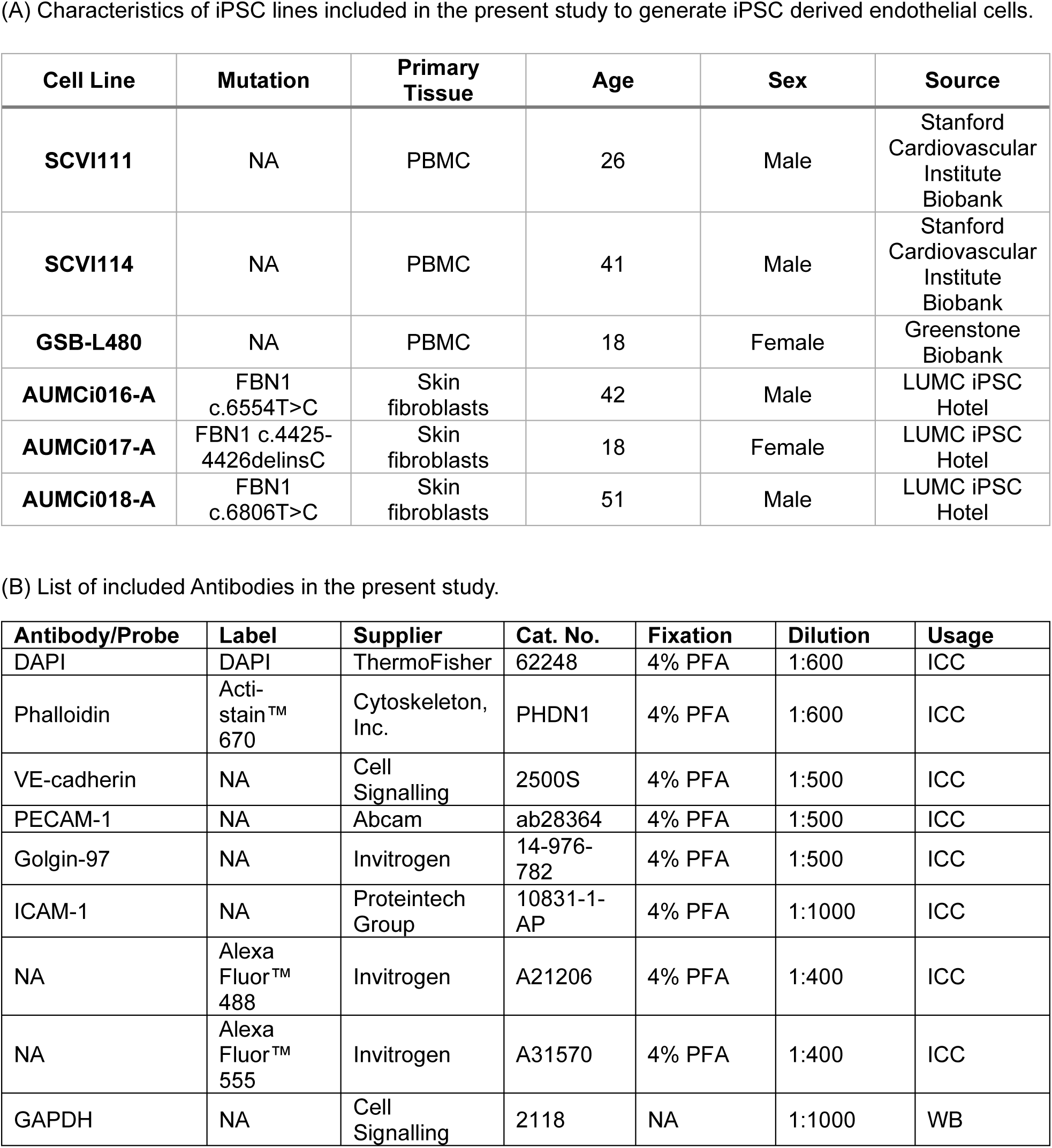
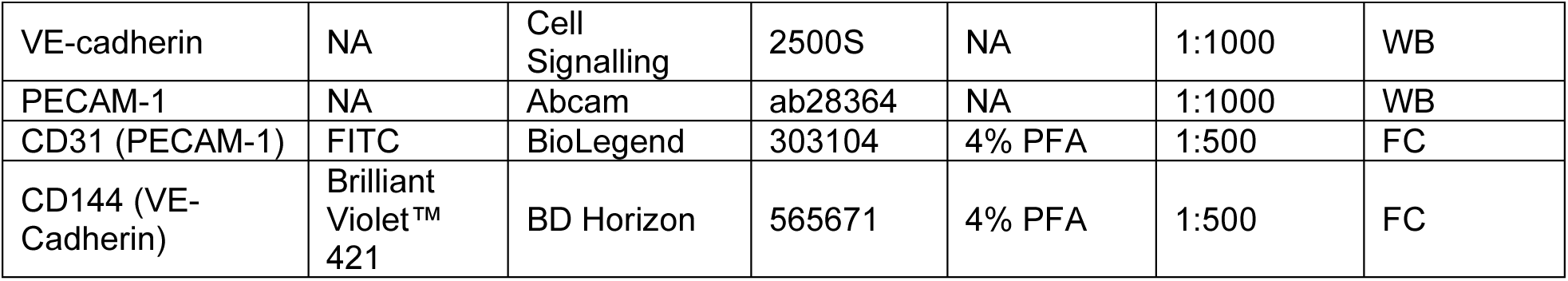
(A) Characteristics of iPSC lines included in the present study to generate iPSC derived endothelial cells. (B) List of included Antibodies in the present study.

## STAR Methods

### hiPSC line culture and maintenance

hiPSCs were cultured on Vitronectin XF (StemCell Technologies) coated plates (Greiner). Media change was performed daily using mTeSR Plus (StemCell Technologies). Cells were passaged as colonies using Gentle Cell Dissociation Reagent according to the manufacturer’s instruction (StemCell Technologies) in a ratio of 1:20. Three independent established hiPSC lines from the Stanford Cardiovascular Institute (SCVI) and Greenstone Biosciences (GSB) biobanks (SCVI-111, SCVI-114 and GSB-L480) were used for control hiPSC-ECs, and were previously reprogrammed from peripheral blood mononuclear cells from healthy individuals. Marfan patient derived cell lines AUMCi016-A, AUMCi017-A, AUMCi018-A were reprogrammed by the LUMC hiPSC hotel using episomal vector expression and p53 knockdown (Okita et al., 2011) and characterized for by our group including pluripotency, trilineage differentiation potential and confirmation of the expected disease-associated variant (Hauger et al., 2026). Information on included cell lines is provided in Table S1 (A). Cells where frozen as clumps in PSC cryopreservation medium (Gibco) and thawed in mTeSR Plus supplemented with 10µM Y-27632 (StemCell Technologies). Master cell banks used for control lines were at passage 26 (SCVI111), passage 41 (SCVI114), passage 18 (GSB-L480) and passage 12 for AUMCi016-A, AUMCi017-A and AUMCi018-A, cells were tested negatively for mycoplasma before use.

### Differentiation of hiPSC-ECs

Human induced pluripotent stem cells (hiPSCs) were differentiated toward an EC phenotype using established protocols with minor adjustments (Orlova et al., 2014). In brief, to start differentiation of hiPSCs, mTeSR Plus was replaced with B(P)EL and mesodermal fate restriction was induced on days 0–3 by supplementing with CHIR99021 (8 μM; Tocris Bioscience). On day 3, endothelial specification was initiated by culturing cells in B(P)EL medium containing vascular endothelial growth factor (VEGF; 50 ng/ml; PeproTech) and the TGF-β pathway inhibitor SB431542 (10 μM; Tocris Bioscience), with medium renewal on days 3, 6, and 9. On day 10, endothelial cells were enriched via magnetic separation using CD31-conjugated Dynabeads (Thermo Fisher Scientific). Purified hiPSC-derived ECs were subsequently expanded in endothelial serum-free medium (EC-SFM; Gibco) supplemented with 1% human platelet-poor serum (Sigma), VEGF (30 ng/ml; PeproTech), and basic fibroblast growth factor (bFGF; 20 ng/ml; Miltenyi Biotec). Cells were cryopreserved at passage 1 using a freezing medium consisting of 40% ECGM-2 (PromoCell), 50% fetal bovine serum (Gibco), and 10% DMSO (Sigma-Aldrich). B(P)EL medium consists of a 1:1 mixture of IMDM (Gibco) and F12 Nutrient Mix (Gibco) supplemented with 5% PFHMII (Gibco), 0.25% BSA (Bovogen Biologicals), 1× chemically defined lipids (Gibco), 0.1× ITS-X (Gibco), 450 μM α-monothioglycerol (Sigma-Aldrich), 0.05 mg/mL ascorbic acid 2-phosphate (Sigma-Aldrich), 2 mM GlutaMAX, (Gibco) and 0.5% penicillin–streptomycin (Gibco).

### Flow Cytometry

Cells were washed with PBS-/- and detached by trypsinization for 5 minutes at 37°C. Subsequently, cells were washed once with PBS-/- and resuspended 1 × 10⁶ cells/mL in blocking buffer (10% BSA in PBS-/-), followed by incubation for 15 minutes at 4°C. Cells then were washed with PBA (0.5% BSA in PBS-/-) and centrifuged (300 × g, 5 minutes, 4°C) prior to staining with fluorochrome-conjugated antibodies against CD31 (FITC) and CD144 (BV421) for 1 hour at 4°C. Following staining, cells were washed once with PBS-/- and fixed with 4% paraformaldehyde for 10 minutes on ice. After three washes with PBA, cells were resuspended in PBS-/- and stored at 4°C until acquisition. Flow cytometric analysis was performed using a BD LSRFortessa (BD Biosciences). Data were analyzed with FlowJo software (BD Life Sciences, Ashland, OR, USA). Stained cell populations were defined relative to an unstained control comprising a pooled sample of all cell lines included in the respective experiment. Doublets were excluded by forward and side scatter pulse geometry gating, and dead cells were excluded based on Live/Dead staining prior to analysis.

### Western Blot

Cells were rinsed with PBS-/- prior to lysis in 2 × SDS sample buffer (125 mM Tris-HCl, pH 6.8, 4% SDS, 20% glycerol, 100 mM DTT, 0.02% bromophenol blue in Milli-Q water). Protein samples were resolved by SDS-PAGE using commercial PROTEAN® TGX™ Precast Protein Gels (Bio-Rad) and subsequently transferred onto nitrocellulose membranes. Membranes were blocked for 1 hour in 5% BSA in TBS-T and incubated with primary antibodies in 5% BSA/TBS-T overnight at 4°C. Following incubation with appropriate secondary antibodies in 5% BSA/TBS-T, signal detection was carried out using ECL Prime Western Blotting Detection Reagent (Amersham/GE Healthcare, Amersham, UK), and chemiluminescent signals were captured with an Amersham AI600 imaging system. Band intensities were quantified by densitometric analysis using ImageQuant TL software (version v8.2.0.0, Cytiva, Marlborough, MA, USA).

### Motility analysis

For quantification of single-cell migration, cells were seeded at low density in 6-well culture plates pre-coated with 0.1% gelatin. Cells were maintained in endothelial serum-free medium (EC-SFM; Gibco) supplemented with 1% human platelet-poor serum (Sigma), VEGF (30 ng/mL; PeproTech), and bFGF (20 ng/mL; Miltenyi Biotec). Plates were transferred to a bench-top fluorescence microscope (BZ-X; Keyence) equipped with environmental control to maintain physiological conditions (37°C, 5% CO₂). Time-lapse imaging was initiated 2h after seeding and performed at 5-minute intervals over a 12-hour period. Single-cell migration was tracked using the Manual Tracking plugin in ImageJ (Fabrice Cordelières, Institut Curie). The resulting cell trajectories were exported and further analyzed using a custom Python script to extract quantitative migration parameters. The directionality index was defined and calculated as the straight-line distance from the first to the last cell position divided by the total distance travelled.

### Endothelial barrier function measurements

Endothelial barrier function was assessed using electric cell–substrate impedance sensing (ECIS) as previously described (C Tiruppathi 1, 1992). In brief, cells were seeded at a density of 125,000 cells/cm² onto fibronectin-coated ECIS arrays (96W10idf; Applied Biophysics, Troy, NY, USA). Transendothelial electrical resistance was continuously monitored at a frequency of 4,000 Hz. Cells were seeded in in endothelial serum-free medium (EC-SFM; Gibco) supplemented with 1% human platelet-poor serum (Sigma), VEGF (30 ng/mL; PeproTech), and bFGF (20 ng/mL; Miltenyi Biotec). After 48 hours a media change was performed to endothelial serum-free medium (EC-SFM; Gibco) supplemented with 1% human platelet-poor serum (Sigma), VEGF (0.5 ng/mL; PeproTech) and bFGF (10 ng/mL; Miltenyi Biotec). Compound treatments were performed 24 hours after the latter media change.

### Immunocytochemistry

Cells were plated onto ibiTreat-treated µ-Slide 8 Well chambers (ibidi GmbH, Germany) coated with 0.1% gelatin. Cells were fixed in 4% paraformaldehyde (Thermo Fisher Scientific) in PBS-/- (Gibco) for 15 minutes at room temperature. After fixation, cells were rinsed three times with PBS-/- and permeabilized using 0.2% Triton X-100 in PBS-/- for 3 minutes. Non-specific binding was blocked by incubation in PBS-/- containing 1% human serum albumin for 1 hour. Primary antibodies were diluted in blocking solution and applied for 1 hour at room temperature, followed by three washes with PBS-/- (see antibody list in Table S1 (B)). Secondary antibody and fluorescent probe incubation followed for 1 hour at room temperature. Subsequently, samples were washed three times with PBS-/-, and stored at 4°C until imaging. Confocal laser scanning microscopy was carried out using a Nikon A1R confocal system (Nikon, Tokyo, Japan). Acquired images were processed and uniformly adjusted using ImageJ (version 1.54d; National Institutes of Health, USA), QuPath (Bankhead et al., 2017), Polarity-JaM (Giese et al., 2025), or custom-developed Python scripts, as indicated for the respective analysis. For quantification of VE-cadherin, ICAM-1, and F-actin mean fluorescence intensity (MFI), cells were segmented using CellPose 2.0 (Pachitariu and Stringer, 2022) integrated in QuPath, and fluorophore-specific MFI was measured within the resulting ROIs.

### ROS measurement assay

Intracellular ROS levels were measured using 5 µM H2DCFDA (Thermo Fisher Scientific). Cells were incubated with H2DCFDA and the respective treatment compounds for 30 min at 37°C in the dark, washed with PBS +/+ with 1 mg/mL D-Glucose (Gibco), and fluorescence was measured using a SpectraMax iD3 plate reader (Ex/Em: 492–495/517– 527 nm). Fluorescence intensities were background-corrected by subtraction of background fluorescence values measured in a control well without H2DCFDA and normalized to cell count per well and the respective PBS-treated controls.

### Flow experiments

To expose hiPSC-derived endothelial cells to laminar shear stress, a computer-controlled perfusion system (ibidi) comprising a pump, fluidic unit, and perfusion set (15 cm tubing, 1.6 mm inner diameter, 10 ml reservoirs) was used. Cells were seeded onto fibronectin-coated µ-Slide VI 0.6 chambers (ibidi) at a concentration of 0.5 × 10⁶ cells/ml and allowed to adhere for 24 hours under static conditions. Subsequently, slides were connected to the perfusion system and subjected to gradually increasing laminar flow, beginning with 2.5 dyn/cm² for 1 hour, followed by 7.5 dyn/cm² for 1 hour, and then maintained at 18 dyn/cm² for 72 hours. All flow experiments were conducted in a standard cell culture incubator at 37 °C and 5% CO₂ using endothelial serum-free medium supplemented with 1% human platelet-poor serum, VEGF (30 ng/ml; PeproTech), and bFGF (20 ng/ml; Miltenyi Biotec).

### Quantification of shear stress parameters

Images were obtained on fixed samples using a Nikon A1R confocal microscope. A maximum intensity projection of 10µm stacks was performed prior to analysis. Cellular detection and feature extraction was then performed using the Polarity-JaM toolbox (Giese et al., 2025) with integrated Cellpose2 algorithm (Pachitariu and Stringer, 2022). Feature quantification was performed by using the web application of Polarity-Jam. Cellular parameters, including nuclei–Golgi orientation, eccentricity and cell orientation, were quantified by calculating the polarity index (PI), which reflects the degree of clustering within a circular distribution and ranges from 0 to 1. Cell eccentricity (elongation; 0 = circular, 1 = elongated), cell shape orientation (alignment relative to the horizontal axis, 0-180°), and organelle orientation (Golgi–nucleus vector, 0-360°).

### RNA sequencing

Total RNA was isolated using the miRNeasy Micro Kit (Qiagen) with on-column DNase digestion (RNase-Free DNase Set, Qiagen) to eliminate genomic DNA contamination. RNA quality and library integrity were assessed using the LabChip Gx Touch system (PerkinElmer). After normalization, 500 ng–1 µg of total RNA was used for library preparation, and sequencing was performed on an Illumina NextSeq500 platform using a P3 flow cell. Raw reads were quality-trimmed using Trimmomatic (v0.39), removing bases below a mean quality score of Q15 within a 5-nucleotide sliding window and retaining reads longer than 15 nucleotides. Filtered reads were aligned to the human reference genome (hg38, Ensembl release 109) using STAR (v2.7.11b). Alignments were processed to remove duplicate reads (Picard v3.1.1), multi-mapping reads, and reads mapping to ribosomal or mitochondrial sequences. Gene-level counts were generated using featureCounts (v2.0.4), considering strand specificity and excluding reads overlapping multiple genes. The resulting count matrix was normalized and differential expression analysis was performed using DESeq2 (v1.36.0). Genes were considered significantly differentially expressed at average count > 5, adjusted p-value < 0.05, and |log2 fold change| > 0.585. Functional annotation was complemented with UniProt data. Downstream analyses were conducted using normalized counts and included volcano and MA plots, hierarchical clustering heatmaps based on Euclidean distance of regularized log-transformed counts, and principal component analysis performed with FactoMineR. Gene set overrepresentation analysis was performed using KOBAS with Benjamini– Hochberg correction (adjusted p < 0.2), including both combined and direction-specific analyses of up- and down-regulated genes. Gene set enrichment analysis (GSEA) was conducted using fGSEA with MSigDB h and c2 collections, ranking genes by signed log10-transformed DESeq2 p-values, and filtering results at adjusted p < 0.2. Transcription factor binding site enrichment analysis was performed using Pscan with CORE vertebrate non-redundant position weight matrices from JASPAR, assessing promoter regions (−450 to +50 bp relative to TSS) of protein-coding genes, with differentially expressed genes as foreground and all genes as background. Statistical significance for differential expression (DESeq2), overrepresentation (KOBAS), and GSEA analyses was determined using Benjamini–Hochberg adjusted p-values, while TFBS enrichment was evaluated using uncorrected p-values as implemented by the respective tools.

## Statistical analysis

Statistical analyses were conducted using GraphPad Prism (GraphPad Software v10.6.0, San Diego, CA, USA). Results are expressed as either mean ± standard error of the mean (SEM) or mean ± standard deviation (SD), as indicated in the figure legends. Data distribution was assessed for normality prior to analysis, and either parametric or non-parametric tests were selected accordingly. Repeated-measures approaches were applied for longitudinal datasets when appropriate. A p-value < 0.05 was considered statistically significant, denoted as *p < 0.05, **p < 0.01, ***p < 0.005, and ****p < 0.001. A limited number of graphical elements used in the figures were adapted from BioRender.com and subsequently modified by the authors using Adobe Illustrator.

## Ethics statement

Human stem cell lines were used in accordance with Amsterdam UMC regulations and institutional oversight procedures, with documented donor informed consent for research and future research. Details about hiPSC lines used for experiments in this work is provided in the STAR methods and in Table S1 (A).

