## supplemental files for "Marfan Patient iPSC-Derived Endothelial Cells Carrying *FBN1* Variants Reveal Endothelial Dysfunction"

**Short title**: Endothelial Dysfunction in Marfan Syndrome hiPSC-Derived Endothelial Cells

**Corresponding authors:**

1) Philipp C. Hauger,

2) Peter L. Hordijk,

3) Vivian de Waard,

Address: Department of Physiology, O2 gebouw, De Boelelaan 1108, 1081 HZ Amsterdam

*Equal contribution

Key words: Marfan, endothelial cells, hiPSCs, disease modeling, inflammation

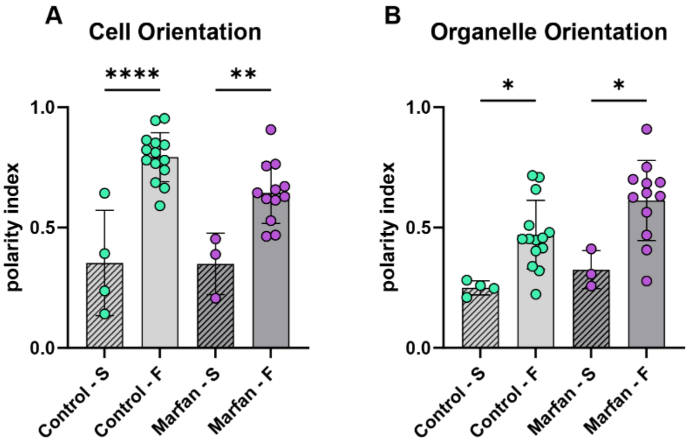


**Supplementary Figure S1: Flow response in iPSC derived ECs.**

Quantification of cell eccentricity (A) and organelle orientation (B) in iECs and iMFS-ECs in static condition and after exposure to LSS. S = Static, F = Flow (LSS). Each datapoint represents one analyzed image, averaged by the Polarity-JaM Web-app from extracted features generated by the analysis pipeline. Statistical analyses were performed on pooled individual datapoints from healthy control and patient samples. Sample numbers per line (LSS): SCVI111 n = 8, SCVI114 n = 6, AUMCi016-A n = 5, AUMCi017-A n = 4, AUMCi018-A n = 3. Sample numbers per line (static): SCVI111 n = 2, SCVI114 n = 2, AUMCi016-A n = 1, AUMCi017-A n = 1, AUMCi018-A n = 1. Data passed the Shapiro–Wilk normality test and groups were compared using a one-way ANOVA. Data are shown as mean ± SD. *$p<0.05$, **$p<0.01$, ****$p<0.0001$, ns $p>0.05$.

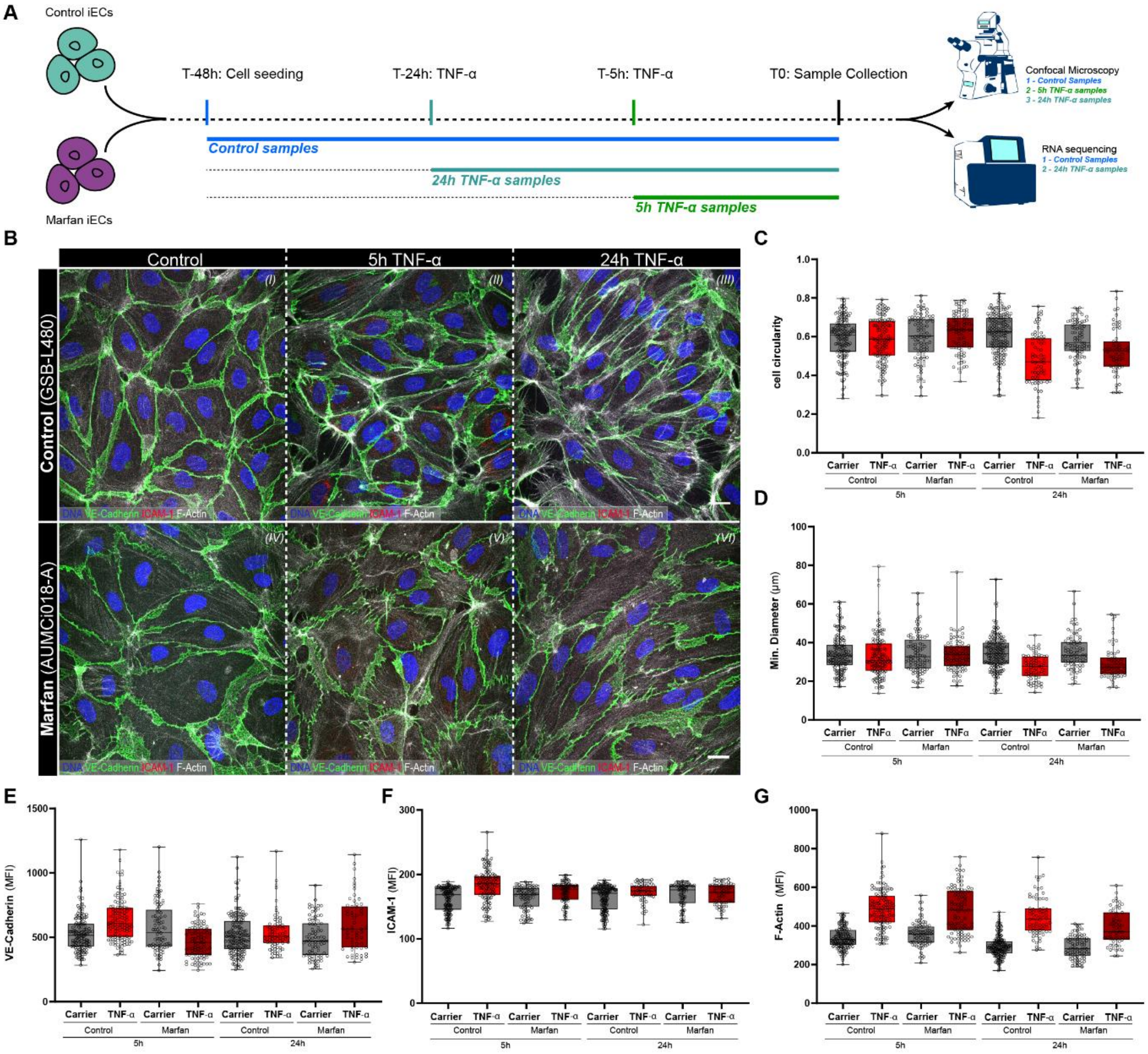


**Supplementary Figure S2: Inflammatory response of iMFS-ECs after TNF-α stimulation.**

(A) Schematic overview of experimental layout to investigate TNF-α response in iECs and iMFS-ECs. Cells were stimulated with 10 ng/mL TNF-α for 5h or 24h, and a vehicle control was established using 24h vehicle stimulation. Cell samples were then analyzed using confocal imaging (24h and 5h time points), as well as RNA sequencing (24h samples). (B) Representative confocal images of iECs (top panel) and iMFS-ECs (bottom panel) after stimulation with vehicle control (I + IV), 5h TNF-α (II + V) or 24h TNF-α (III + VI), blue: DNA, green: VE-Cadherin, red: ICAM1, gray: F-Actin. Scale bar: 20µm. (C–G) Single-cell quantification of confocal images using QuPath. Three images per iEC line (SCVI111, GSB-L480) and per iMFS-EC line (AUMCi016-A, AUMCi017-A) were acquired for each experimental condition. Datapoints represent individual cells and are shown as box plots with whiskers indicating minimum to maximum values. (C) Cell circularity (values between 0 and 1; 1 = circular). (D) Minimal cellular diameter (µm). (E) VE-cadherin mean fluorescence intensity (MFI). (F) ICAM-1 MFI. (G) F-actin MFI.

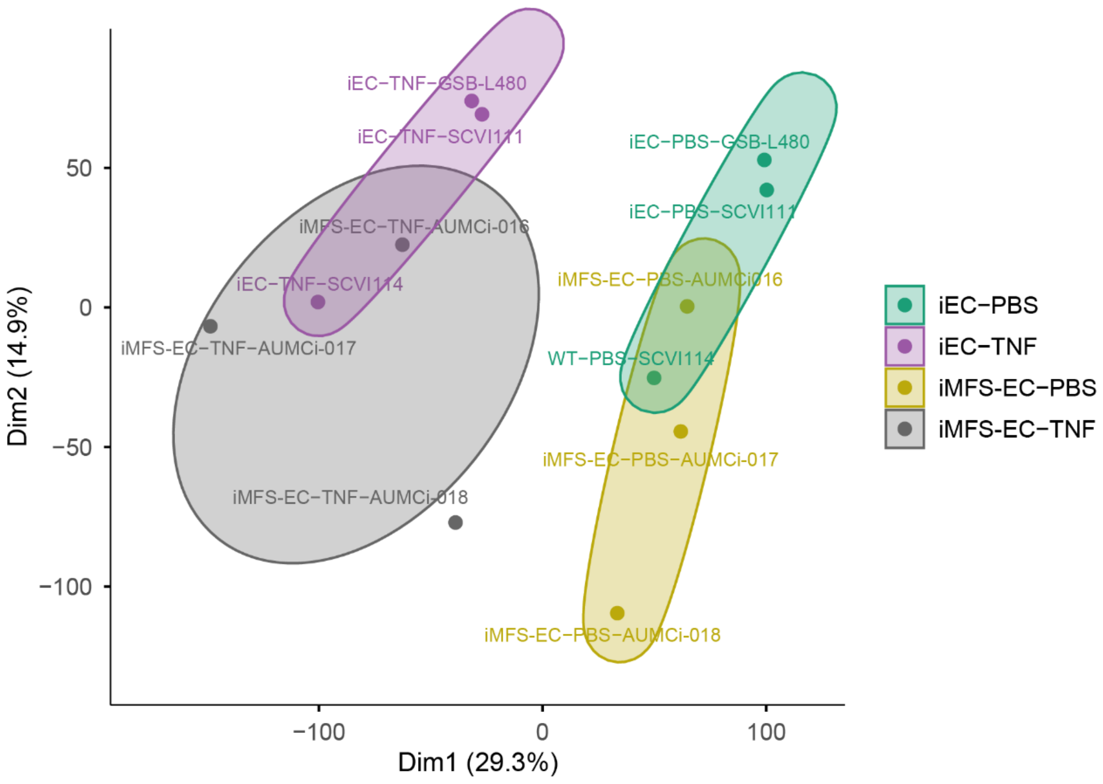


**Supplementary Figure S3: PCA plot.**

Principal component analysis (PCA) of bulk RNA-seq samples. PCA was performed on regularized log-transformed normalized gene counts generated with DESeq2. Each point represents an individual sample, colored according to experimental condition. Principal components 1 and 2 explain 29.3% and 14.9% of the total variance, respectively.

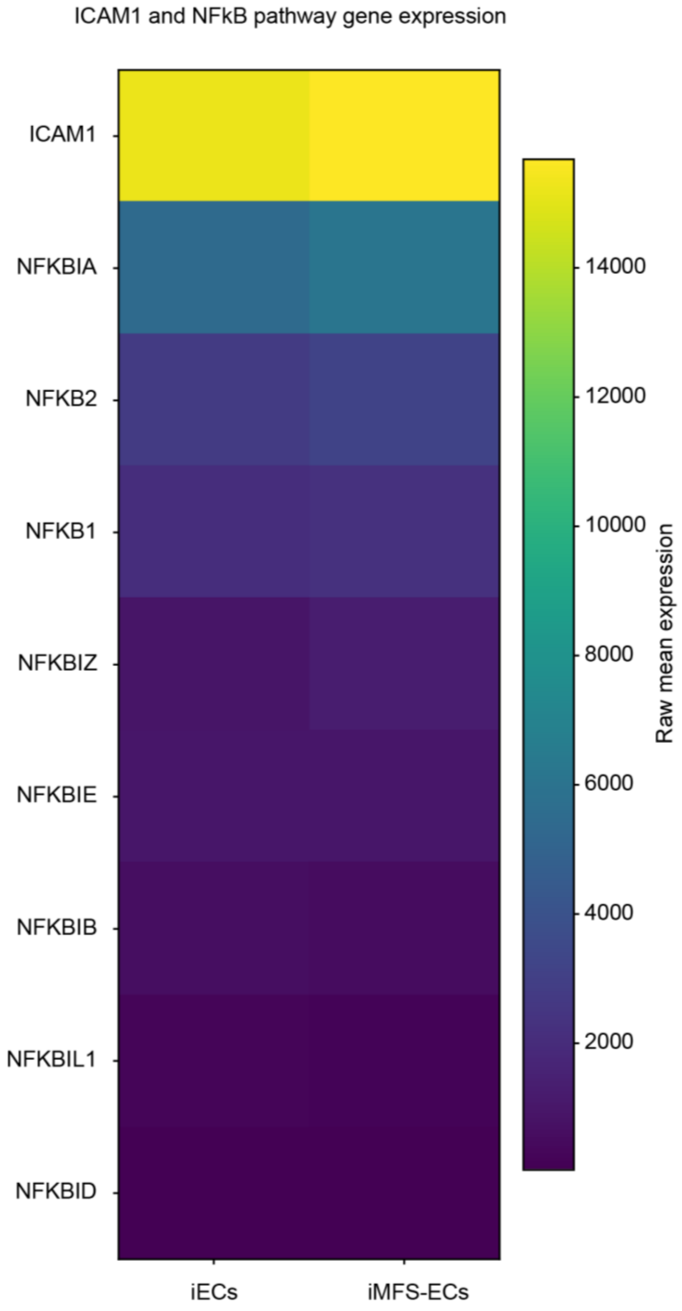


**Supplementary Figure S4: Heatmap for inflammatory gene expression.**

Heatmap visualizing gene expression plotting total reads of ICAM1 and NF-κB pathway genes for $iEC_{TNF\text{-}\alpha}$ versus $iMFS\text{-}EC_{TNF\text{-}\alpha}$ comparison. $Padj_{iEC\text{-}iMFS\text{-}EC}$ for all genes >0.05.

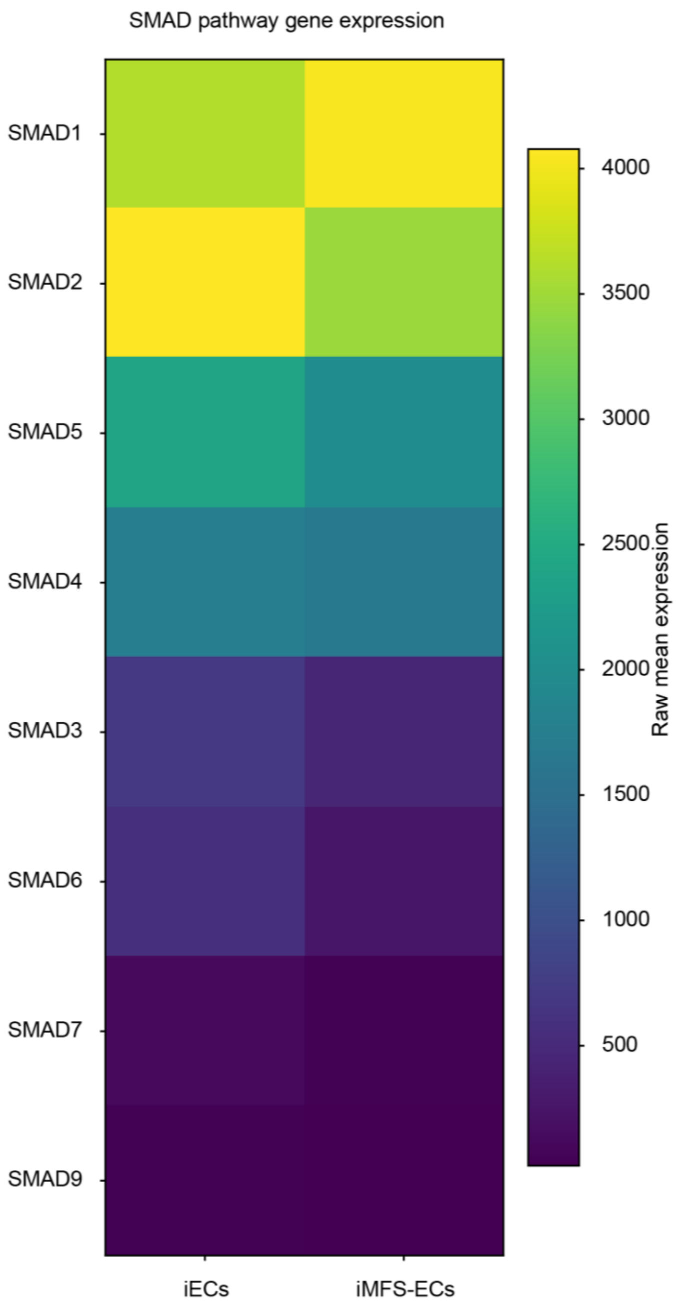


**Supplementary Figure S5: Heatmap for SMAD pathway.**

Heatmap visualizing gene expression plotting total reads of genes in the SMAD pathway for $iEC_{Vehicle\text{-}Control}$ versus $iMFS\text{-}EC_{Vehicle\text{-}Control}$ comparison. $Padj_{iEC\text{-}iMFS\text{-}EC}$ for all genes >0.05.

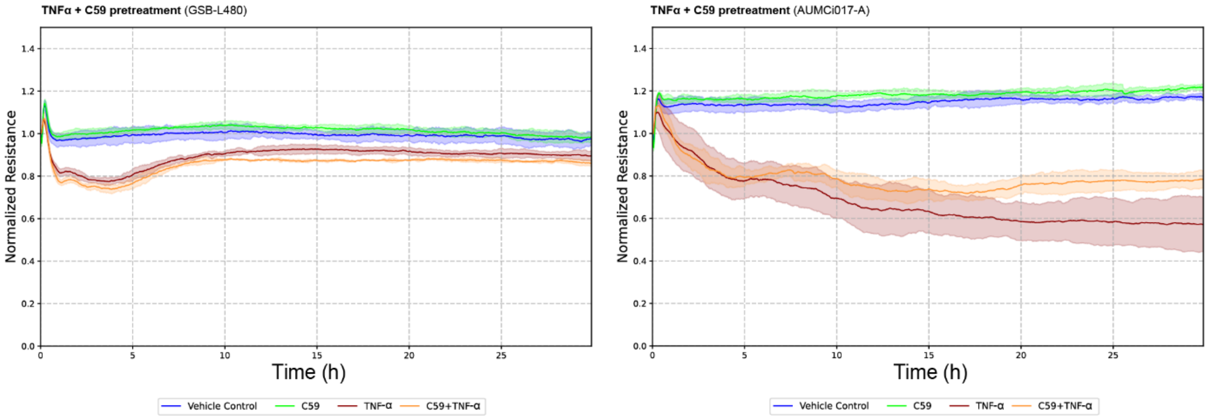


**Supplementary Figure S6: C59 pretreatment.**

Representative ECIS graphs showing EC response of (A) GSB-L480 and (B) AUMCi16-A to 10 µg/mL TNF-α (red), to 10 µg/mL TNF-α following 2h pretreatment with 2 µM C59 (orange), to 2µM C59 (green) or a vehicle control (blue). Each data point represents the normalized resistance to a respective vehicle control per cell line and time point of treatment.

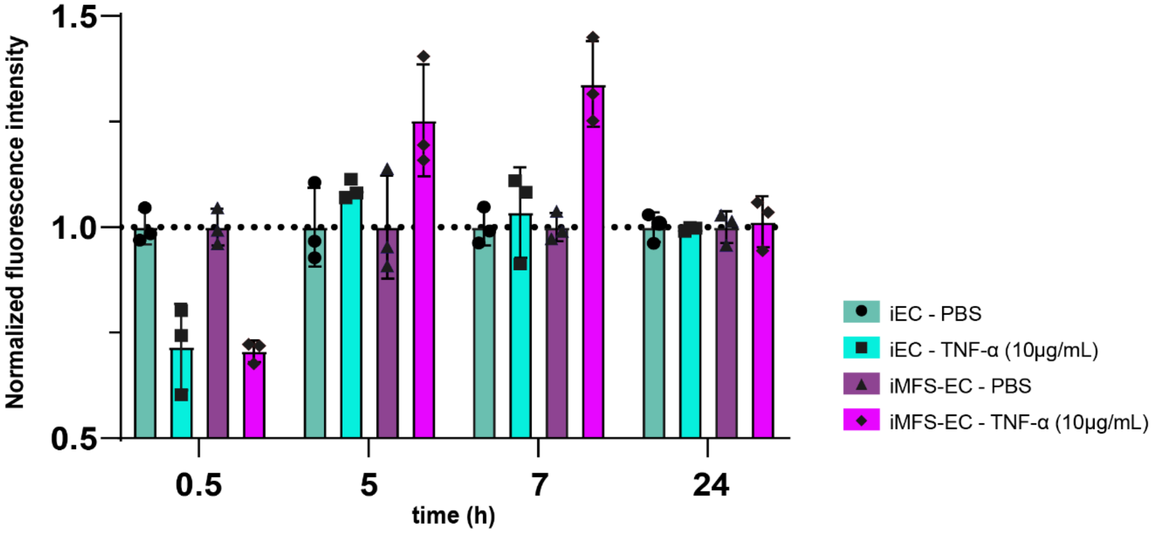


**Supplementary Figure S7: ROS measurements.**

Analysis of ROS levels after stimulation with 10ng/mL TNF-α or a carrier control (PBS-/-) for 0.5h, 5h, 7h and 24h by measuring H2DCFDA fluorescence intensity. Fluorescence was measured in triplicates using a 96-well plate reader. Raw fluorescence values were background-corrected by subtraction of background fluorescence values measured in a control well and normalized to nucleus-based cell count for each well. Data represent triplicate measurements normalized to the respective PBS control.

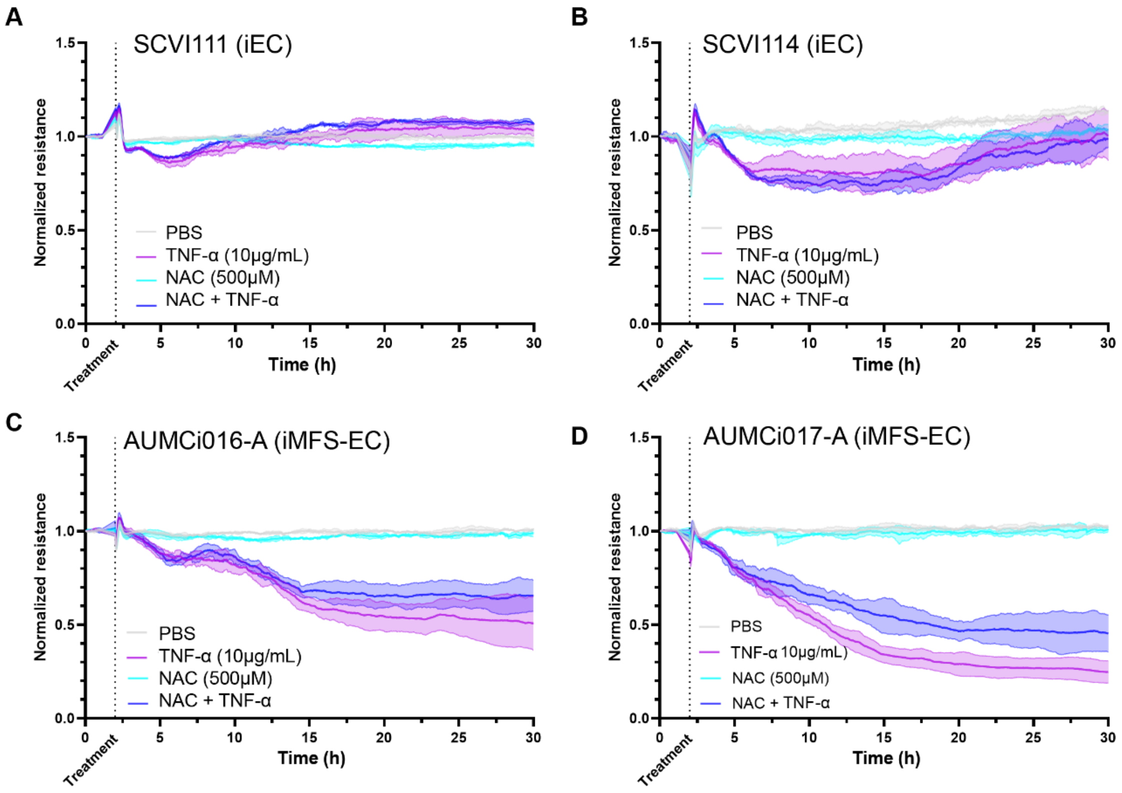


**Supplementary Figure S8: NAC pretreatment.**

Representative ECIS graphs showing EC response of (A) SCVI111, (B) SCVI114, (C) AUMCi16-A, (D) AUMCi017-A to 10 ng/mL TNF-α (pink), 500µM n-Acetylcysteine (cyan) or combined treatment with 10 ng/mL TNF-α and 500µM n-Acetylcysteine (blue). Carrier control treatment corresponding to the combined treatment is shown in gray. Each data point represents the normalized resistance to a respective vehicle control per cell line and time point of treatment.

**Table S1:**

(A) Characteristics of iPSC lines included in the present study to generate iPSC derived endothelial cells.

| Cell Line | Mutation | Primary Tissue | Age | Sex | Source |
|---|---|---|---|---|---|
| **SCVI111** | NA | PBMC | 26 | Male | Stanford Cardiovascular Institute Biobank |
| **SCVI114** | NA | PBMC | 41 | Male | Stanford Cardiovascular Institute Biobank |
| **GSB-L480** | NA | PBMC | 18 | Female | Greenstone Biobank |
| **AUMCi016-A** | FBN1 c.6554T>C | Skin fibroblasts | 42 | Male | LUMC iPSC Hotel |
| **AUMCi017-A** | FBN1 c.4425-4426delinsC | Skin fibroblasts | 18 | Female | LUMC iPSC Hotel |
| **AUMCi018-A** | FBN1 c.6806T>C | Skin fibroblasts | 51 | Male | LUMC iPSC Hotel |

(B) List of included Antibodies in the present study.

| Antibody/Probe | Label | Supplier | Cat. No. | Fixation | Dilution | Usage |
|---|---|---|---|---|---|---|
| DAPI | DAPI | ThermoFisher | 62248 | 4% PFA | 1:600 | ICC |
| Phalloidin | Acti-stain™ 670 | Cytoskeleton, Inc. | PHDN1 | 4% PFA | 1:600 | ICC |
| VE-cadherin | NA | Cell Signalling | 2500S | 4% PFA | 1:500 | ICC |
| PECAM-1 | NA | Abcam | ab28364 | 4% PFA | 1:500 | ICC |
| Golgin-97 | NA | Invitrogen | 14-976-782 | 4% PFA | 1:500 | ICC |
| ICAM-1 | NA | Proteintech Group | 10831-1-AP | 4% PFA | 1:1000 | ICC |
| NA | Alexa Fluor™ 488 | Invitrogen | A21206 | 4% PFA | 1:400 | ICC |
| NA | Alexa Fluor™ 555 | Invitrogen | A31570 | 4% PFA | 1:400 | ICC |
| GAPDH | NA | Cell Signalling | 2118 | NA | 1:1000 | WB |
| VE-cadherin | NA | Cell Signalling | 2500S | NA | 1:1000 | WB |
| PECAM-1 | NA | Abcam | ab28364 | NA | 1:1000 | WB |
| CD31 (PECAM-1) | FITC | BioLegend | 303104 | 4% PFA | 1:500 | FC |
| CD144 (VE-Cadherin) | Brilliant Violet™ 421 | BD Horizon | 565671 | 4% PFA | 1:500 | FC |

**STAR methods**

### Key resources table

| REAGENT or RESOURCE | SOURCE | IDENTIFIER |
|---|---|---|
| Antibodies | | |
| VE-cadherin | Cell Signalling | Cat#2500S; RRID: AB_10839118 |
| PECAM-1 | Abcam | Cat#ab28364; RRID: AB_726362 |
| Golgin-97 | Invitrogen | Cat#14-9767-82; RRID: AB_2573010 |
| ICAM-1 | Proteintech Group | Cat#10831-1-AP, RRID: AB_2264494 |
| Alexa Fluor™ 488 | Invitrogen | Cat#A21206; RRID: AB_2535792 |
| Alexa Fluor™ 555 | Invitrogen | Cat#A31570; RRID: AB_2536180 |
| GAPDH | Cell Signalling | Cat#2118; RRID: AB_561053 |
| CD31 (PECAM-1) | BioLegend | Cat#303104; RRID: AB_314330 |
| CD144 (VE-Cadherin) | BD Horizon | Cat#565671; RRID: AB_2744284 |
| Chemicals, peptides, and recombinant proteins | | |
| Vitronectin XF™ | StemCell Technologies | Cat#100-0763 |
| Gentle Cell Dissociation Reagent | StemCell Technologies | Cat#100-0485 |
| Y-27632 | StemCell Technologies | Cat#72302 |
| CHIR99021 | Tocris Bioscience | Cat#4423 |
| VEGF | ThermoFisher (PeproTech) | Cat#100-20-100UG |
| SB431542 | Tocris Bioscience | Cat#1614 |
| Dynabeads™ CD31 Endothelial Cell | ThermoFisher (Invitrogen) | Cat#11155D |
| Human Endothelial SFM | ThermoFisher (Gibco) | Cat#11111044 |
| human platelet-poor serum | Sigma-Aldrich | Cat#P2918-20ML |
| Human FGF-2 (bFGF) | Miltenyi Biotec | Cat#130-093-838 |
| Endothelial Cell Growth Medium 2 Kit (ECGM-2) | Promocell | Cat#C-22111 |
| H2DCFDA | ThermoFisher (Invitrogen) | Cat#D399 |
| IMDM, no phenol red | ThermoFisher (Gibco) | Cat#21056023 |
| Ham's F-12 Nutrient Mix | ThermoFisher (Gibco) | Cat#11765054 |
| PFHM-II Protein-Free Hybridoma Medium (liquid) | ThermoFisher (Gibco) | Cat#12040077 |
| Bovine Serum Albumin | Bovogen Biologicals | Cat#BSAS-AU |
| Chemically Defined Lipid Concentrate | ThermoFisher (Gibco) | Cat#11905031 |
| Insulin-Transferrin-Selenium-Ethanolamine (ITS -X) | ThermoFisher (Gibco) | Cat#51500056 |
| 1-Thioglycerol (αMTG) | Sigma-Aldrich | Cat#M6145-100ML |
| L-Ascorbic acid 2-phosphate sesquimagnesium salt hydrate (AA2P) | Sigma-Aldrich | Cat#A8960-5G |
| GlutaMAX™ Supplement | ThermoFisher (Gibco) | Cat#35050061 |

| | | |
|---|---|---|
| Penicillin-Streptomycin | ThermoFisher (Gibco) | Cat#15140122 |
| DAPI solution | ThermoFisher (Invitrogen) | Cat#62248 |
| DPBS (1X) | ThermoFisher (Gibco) | Cat#14190144 |
| C59 | Selleckchem | Cat#S737 |
| N-Acetyl-L-cysteine | Sigma-Aldrich | A7250-5G |
| Recombinant Human TNF-alpha | PeproTech | Cat#300-01A-10UG |
| Human IL-1ß | PeproTech | 200-01B-10UG |
| TFLLR-NH2 trifluoroacetate salt | Sigma Aldrich | T7830-5MG |
| 8-pCPT-2'-O-Me-cAMP | Biolog | Cat#C 041 |
| Critical commercial assays | | |
| PSC Cryopreservation kit | Thermo Fisher | Cat#A2644601 |
| mTeSR™ Plus | StemCell Technologies | Cat#100-0276 |
| Deposited data | | |
| Bulk RNA sequencing data | This paper | GEO: GSE335638 |
| Experimental models: Cell lines | | |
| hiPSC line | Stanford Cardiovascular Institute Biobank | SCVI111 |
| hiPSC line | Stanford Cardiovascular Institute Biobank | SCVI114 |
| hiPSC line | Greenstone Biobank | GSB-L480 |
| hiPSC line | LUMC iPSC Hotel | AUMCi016-A |
| hiPSC line | LUMC iPSC Hotel | AUMCi017-A |
| hiPSC line | LUMC iPSC Hotel | AUMCi018-A |
| Experimental models: Organisms/strains | | |
| Oligonucleotides | | |
| Recombinant DNA | | |
| Software and algorithms | | |
| Python 3.11.7 | Python Software Foundation | https://www.python.org/ |
| QuPath 0.5.1 | Bankhead et al., 2017 | https://github.com/qupath/qupath |
| Fiji-ImageJ | Shindelin et al., 2012 | https://imagej.net/software/fiji/ |
| GraphPad Prism 9 | GraphPad Software | https://www.graphpad.com/ |
| BioRender | Science Suite Inc. | https://www.biorender.com/ |
| Polarity-JaM | Giese et al., 2025 | https://polarityjam.readthedocs.io/en/latest/ |
| CellPose 2.0 | Pachitariu et al., 2022 | https://www.cellpose.org/ |
| Other | | |

**Resource availability**

**Lead contact**

Requests for additional information, resources, or reagents should be directed to the corresponding authors P. C. Hauger, V. de Waard. or P.Hordijk..
